# The N-terminus of Stag1 is required to repress the 2C program by maintaining rRNA expression and nucleolar integrity

**DOI:** 10.1101/2021.02.14.429938

**Authors:** Dubravka Pezic, Samuel Weeks, Wazeer Varsally, Pooran S. Dewari, Steven Pollard, Miguel R. Branco, Suzana Hadjur

## Abstract

Several studies have shown a role for Stag proteins in cell identity. Our understanding of how Stag proteins contribute to cell identity have largely been focused on its roles in chromosome topology as part of the cohesin complex and the impact on protein-coding gene expression. Furthermore, several Stag paralogs exist in mammalian cells with non-reciprocal chromosome structure and cohesion functions. Why cells have so many Stag proteins and what specific functions each Stag protein performs to support a given cell state are poorly understood. Here we reveal that Stag1 is the dominant paralog in mouse embryonic stem cells (mESC) and is required for pluripotency. Through the discovery of diverse, naturally occurring Stag1 isoforms in mESCs, we shed new light not only on the unique ends of Stag1 but also the critical role that their levels play in stem cell identity. Furthermore, we revel a new role for Stag1, and specifically its unique N-terminal end, in regulating nucleolar integrity and safeguarding mESCs from totipotency. Stag1 is localised to repressive perinucleolar regions, bound at repeats and interacts with Nucleolin and TRIM28. Loss of the Stag1 N-terminus, leads to decreased LINE-1 and rRNA expression and disruption of nucleolar structure and function which consequently leads to activation of the two-cell-like (2C-LC)-specific transcription factor DUX and conversion of pluripotent mESCs to totipotent 2C-LCs. Our results move beyond protein-coding gene regulation via chromatin loops into a new role for Stag1 in repeat regulation and nucleolar structure, and offer fresh perspectives on how Stag proteins contribute to cell identity and disease.

## INTRODUCTION

Cohesin is a ubiquitously expressed, multi-subunit protein complex that has fundamental roles in cell biology including sister chromosome cohesion, 3D chromatin topology and regulation of cell identity ^1-6^. Much of our understanding of how cohesin contributes to cell identity has been studied in the context of its roles in protein-coding gene expression and 3D organization of interphase chromatin structure ^7-15^. Indeed, loss of cohesin and its regulators results in a dramatic loss of chromatin topology at the level of Topologically Associated Domains (TAD) and chromatin loops, albeit with modest changes to gene expression ^16-22^. This suggests that cohesin’s roles in development and disease extend beyond gene expression regulation and highlight the need to re-evaluate how cohesin regulators shape the structure and function of the genome.

The association of cohesin with chromosomes is tightly controlled by several regulators, including the Stromalin Antigen protein (known as Stag or SA), which has been implicated in cell identity regulation and disease development ^2,3,23-26^. Stag proteins interact with the Rad21 subunit of cohesin and mediate its association with DNA and CTCF ^27-30^. Mammalian cells express multiple Stag paralogs, which have >90% sequence conservation in their central domain yet perform distinct functions ^31-34^. It is likely that the divergent N- and C-terminal regions provide functional specificity. For example, the N-terminus of Stag1 contains a unique AT-hook ^35^ which is required for its preferential participation in telomere cohesion ^31^. However, the underlying mechanisms by which Stag proteins and their divergent ends influence cell identity are largely unknown.

The nucleolus is a multifunctional nuclear compartment which coordinates ribosome biogenesis with cell cycle control and mRNA processing ^36^. It forms through self-organization of its constituent proteins and the rDNA gene clusters into a tripartite, phase separated condensate ^37,38^ which is intimately connected to overall nuclear organization ^39^. In line with its liquid-like properties, the nucleolus is itself plastic, undergoing dramatic changes in response to cell cycle, metabolic or developmental cues. For example, functional nucleoli play an important role in the control of cell identity during early mouse development ^40^. Two-cell (2C) stage totipotent embryos exhibit ‘immature’ nucleoli with poorly defined structure and low levels of perinucleolar heterochromatin ^41,42^. This global chromatin accessibility contributes to the expression of the 2C-specific transcription factor DUX and the subsequent activation of MERVL elements ^43,44^. As the embryo reaches the 8-cell stage, cells harbour fully mature phase-separated nucleoli, defined heterochromatin around the nucleolar periphery ^45^ and robust rRNA expression, all of which are essential for cells to commit to differentiation ^40,46^. In contrast, mouse embryonic stem cells (mESC) exhibiting nucleolar stress lead to conversion to 2C-like cell (2C-LC) identity *in vitro* ^47^ and nucleolar proteins that control rRNA transcription and processing are essential for 2C-LC repression ^48^), highlighting the tight relationship between rRNA levels, nucleolar structure and cell identity.

It is known that cohesin is necessary for nucleolar integrity in yeast. Core cohesin subunits have been shown to bind to the non-transcribed region of the rDNA locus ^49^ and the 35S and 5S genes form loops that are dependent on Eco1, the cohesin subunit known to acetylate Smc3 and thus stabilize cohesin rings on chromatin ^50^. Consequently, yeast with eco1 mutations exhibit disorganised nucleolar structure and defective ribosome biogenesis.

Here we reveal a novel role for Stag1, and in particular its unique N-terminal end, in regulating nucleolar integrity and 2C repression to maintain mESC cell identity. Stag1 binds to repeats associated with nucleolar structure and function including rDNA and LINE-1 and interacts with the Nucleolin/TRIM28 complex that resides within perinucleolar chromatin to maintain nucleolar integrity. Loss of Stag1 or specifically the N-terminus in mESCs leads to reduced nascent rRNA and LINE-1, nucleolar disruption, increased expression of DUX and conversion of mESCs to totipotent 2C-LC cells. In addition to presenting a new role for Stag1 in repeat regulation, nucleolar structure and translation control, our results also reveal a previously unappreciated transcriptional diversity of Stag1 in stem cells and highlights the complexity of cohesin regulation in mammalian cells. We show that cells change both the levels of Stag paralogs as well as the balance of isoforms to control cell identity and point to the importance of the divergent, unstructured ends of Stag1 proteins in nuclear body structure and cell fate control. Our results offer fresh perspectives on how Stag proteins, known to be pan-cancer targets ^3^ contribute to cell identity and disease.

## RESULTS

### A functional change in cohesin regulation in cells of different potential

We analysed the expression levels of cohesin regulators in mESCs by qRT-PCR at different stages of pluripotency. During the transition between naïve (2i mESC) and primed epiblast-like (EpiLC) pluripotent cells *in vitro*, levels of the core cohesin subunits Smc1 and Smc3 do not change, while Stag1 becomes downregulated and Stag2 becomes upregulated (Fig. 1a, b, S1a, b). This is supported by western blot (WB) analysis where we observe a 2-3-fold higher level of chromatin-associated Stag1 compared to Stag2 protein in naïve (2i) mESC, while Stag2 levels are 5-10-fold higher in EpiLC (Fig. 1b, S1c). These results, together with similar observations ^26^, identify Stag1 as the dominant paralog in naïve mESC and suggest that a switch between Stag1 and Stag2 may represent a functionally important change in cohesin regulation at different stages of pluripotency.

**Figure 1.**
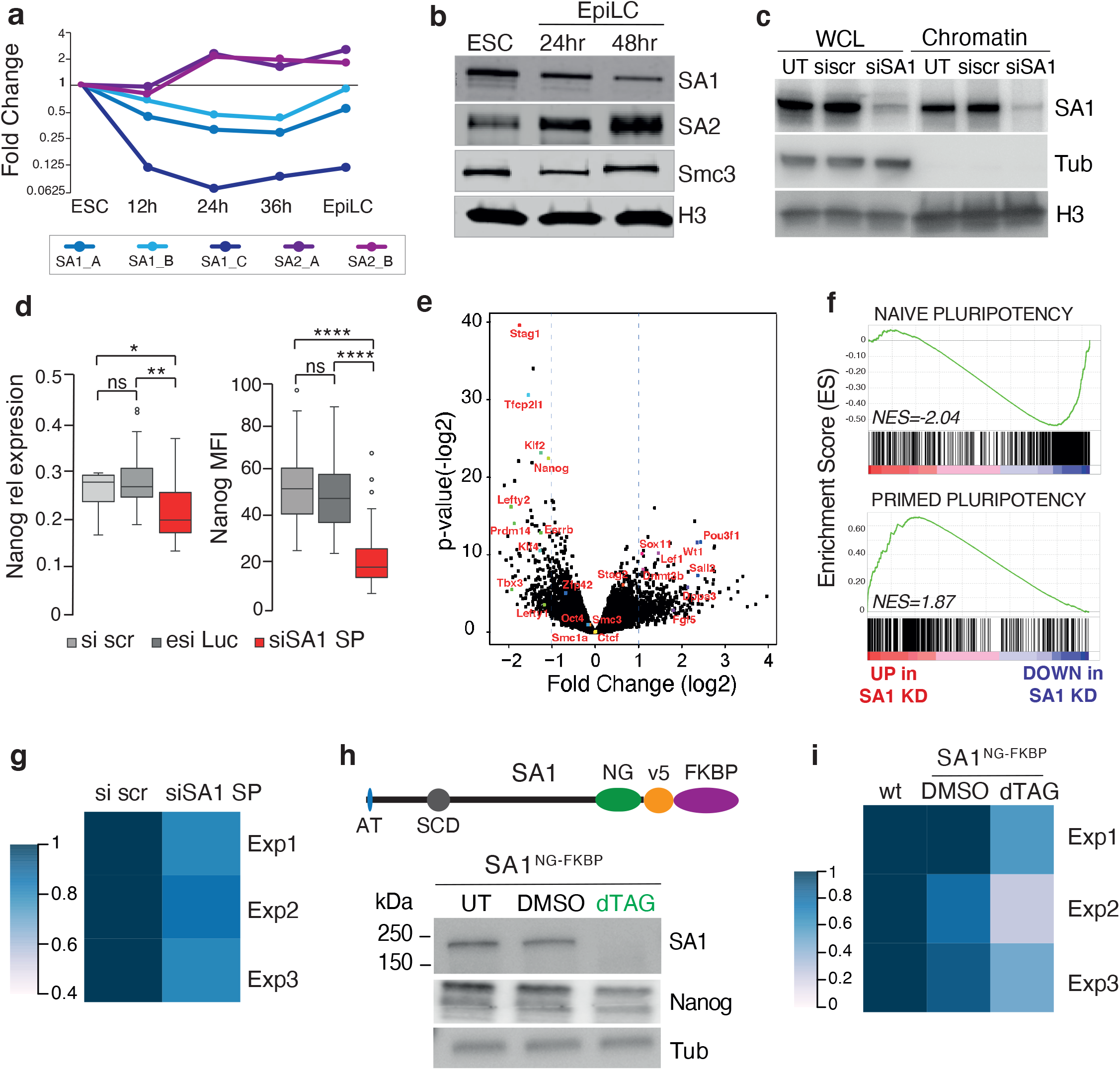
STAG1 is required for naïve pluripotency in mouse ESCs. a) Log2 fold change of Stag1 (SA1) and Stag (SA2) gene expression assessed by qRT-PCR during *in vitro* mESC cell differentiation towards EpiLC. Multiple primer pairs were used for SA1 (blue) and SA2 (purple) mRNA (see box). Data are derived from two biological replicates. b) Whole cell protein extracts (WCL) from naïve mESC and EpiLCs and analysed by western blot (WB) for levels of SA1, SA2 and Smc3. H3 serves as a loading control. c) WB analysis of SA1 levels in WCL and chromatin fractions upon treatment with scrambled control siRNAs (si scr) or SmartPool SA1 siRNAs (siSA1) for 24hr in naïve mESC cells. Tubulin (Tub) and H3 serve as fractionation and loading controls. d) Left, relative expression of Nanog mRNA by qRT-PCR in naïve mESCs upon treatment with si scr, esiLuciferase control or siSA1. Data are from 8 biological replicates. Right, Mean fluorescence intensity (MFI) of Nanog protein assessed by Immunofluorescence (IF) in naïve mESCs treated with same siRNAs as before. Cells were counterstained with DAPI. Data is n>100 cells/condition across 2 biological replicates. Whiskers and boxes indicate all and 50% of values, respectively. Central line represents the median. Asterisks indicate a statistically significant difference as assessed using two-tailed t-test. * p<0.05, ** p<0.005, *** p<0.0005, **** p<0.0001, ns = not significant. e) Volcano plot displaying the statistical significance (-log2 p-value) versus magnitude of change (log2 fold change) from RNA-sequencing data produced in mESCs treated with siscr or siSA1 for 24hrs. Data is from 3 biological replicates. Vertical blue dashed lines represent changes of 2-fold. Selected genes associated with cohesin, pluripotency and differentiation have been highlighted in red. f) Enrichment score (ES) plots from Gene Set Enrichment analysis (GSEA) using curated naïve or primed pluripotency gene sets (see Methods). Negative and positive normalized (NES) enrichment scores point to the gene set being over-represented in the top-most down- or up-regulated genes in SA1 KD mESC, respectively. Vertical bars refer to individual genes in the gene set and their position reflects the contribution of each gene to the NES. g) Area occupied by AP+ colonies in mESCs treated with si scr and si SA1 from three independent biological replicates where n>50 colonies/condition were counted. h) CRISPR/Cas9 was used to knock-in a NeonGreen-v5-FKBP tag on both alleles of endogenous Stag1 at the C-terminus (SA1^NG-FKBP^). The resultant Stag1 protein is 42kDa larger. Shown also are known features of SA1 including the N-terminal AT-hook (AT) and the stromalin conserved domain (SCD). WB analysis of SA1and Nanog levels in a targeted mESC clone after treatment with DMSO or dTAG. Tubulin (Tub) serves as a loading control. i) Analysis of the area occupied by AP+ colonies as above but in WT or SA1^NG-FKBP^ mESC treated with DMSO or dTAG. Data is from three independent biological replicates where n>50 colonies/condition were counted.

### Stag1 is required for pluripotency

To investigate the functional importance of Stag1 in the regulation of pluripotency, we first established a Stag1 knockdown (KD, ‘siSA1’, Methods) strategy using siRNAs. This resulted in a significant reduction of Stag1 at the mRNA and protein levels (4-5-, 8-10-fold, respectively), in both serum-grown (FCS) and naive mESC without affecting the cell cycle (Fig. 1c, S1d-f). Using Nanog as a marker of naïve pluripotency, we observed a significant downregulation of Nanog mRNA and protein levels within 24hrs of Stag1 KD in mESC (Fig. 1d, S1g), suggesting that Stag1 may be required for pluripotency. Global analysis of the mESC transcriptome using RNA-sequencing upon siRNA-mediated Stag1 KD revealed that 375 genes were up- and 205 genes were down-regulated by at least 2-fold (Fig. 1e). Among the downregulated group were genes known to have roles in the maintenance of pluripotency (ie. Nanog, Tbx3, Esrrb, Klf4), while genes associated with exit from pluripotency (Dnmt3b, Fgf5) and differentiation (ie. Pou3f1 (Oct6), Sox11) were upregulated (Fig. 1e). Gene Set Enrichment Analysis (GSEA) ^51,52^ confirmed a reproducible loss of naïve pluripotency-associated gene signature and enrichment for genes associated with primed pluripotency upon Stag1 KD (Fig. 1f, S1h).

The loss of the naïve transcriptional programme upon Stag1 KD suggests that mESCs may require Stag1 for the maintenance of self-renewal. To test this, we plated cells in self-renewal conditions at clonal density and determined the proportion of undifferentiated cells upon Stag1 KD by measuring the area occupied by the colonies with high alkaline phosphatase activity (AP+). In scrambled siRNA-treated controls, 52% of plated cells retain their naïve state, identified by AP+ colonies which was not significantly different from untreated cells. Upon Stag1 KD, both the proportion of AP+ colonies and the area they occupy decreased by an average of 20% compared to siRNA controls, indicating that mESCs have a reduced ability to self-renewal in the absence of Stag1 (Fig. 1g, S5d).

We validated these observations by using CRISPR/Cas9 to knock-in an mNeonGreen-FKBP12^F36V^ tag ^53^ at the C-terminus of both alleles of the endogenous Stag1 locus (SA1^NG_FKBP^) in mESC (Fig. 1h, S1i-k). Upon dTAG addition, Stag1 protein is robustly degraded in a SA1^NG_FKBP^ mESC clone (Fig. 1h, S1k). As we had previously observed with siRNA treatment, dTAG-mediated degradation of Stag1 led to a reduction in Nanog protein (reduced by 24% compared to DMSO controls) (Fig. 1h) and self-renewal potential was reduced by an average of 38% compared to DMSO-treated cells (Fig. 1i). Together, our results are consistent with a requirement for Stag1 in the control of naïve pluripotency.

### STAG1 localizes to both euchromatin and heterochromatin

To understand how Stag1 contributes to pluripotency, we first investigated its subcellular localization. Live-cell imaging of Hoechst-labelled SA1^NG_FKBP^ mESC revealed the expected and predominant localisation of Stag1 in the nucleus with a notable punctate pattern within the nucleoplasm (Fig. 2a). Stag1 was also colocalised with Hoechst-dense regions (Fig. 2a, arrows) and enriched in Hoechst-dense foci compared to the whole nucleus (Fig. 2b). This was of interest since Hoechst stains AT-rich heterochromatin which is enriched around the nucleolus, at the nuclear periphery and in discreet foci within the nucleoplasm ^39,54^. Acute degradation of Stag1 in SA1^NG_FKBP^ mESCs resulted in increased Hoechst signal intensity (Fig. 2c) and a significant increase in Hoechst foci volume (Fig. 2d). siRNA-mediated Stag1 KD mESCs revealed similar changes to heterochromatin, as assessed by DAPI and H3K9me3 staining (Fig. S2a, b).

**Figure 2.**
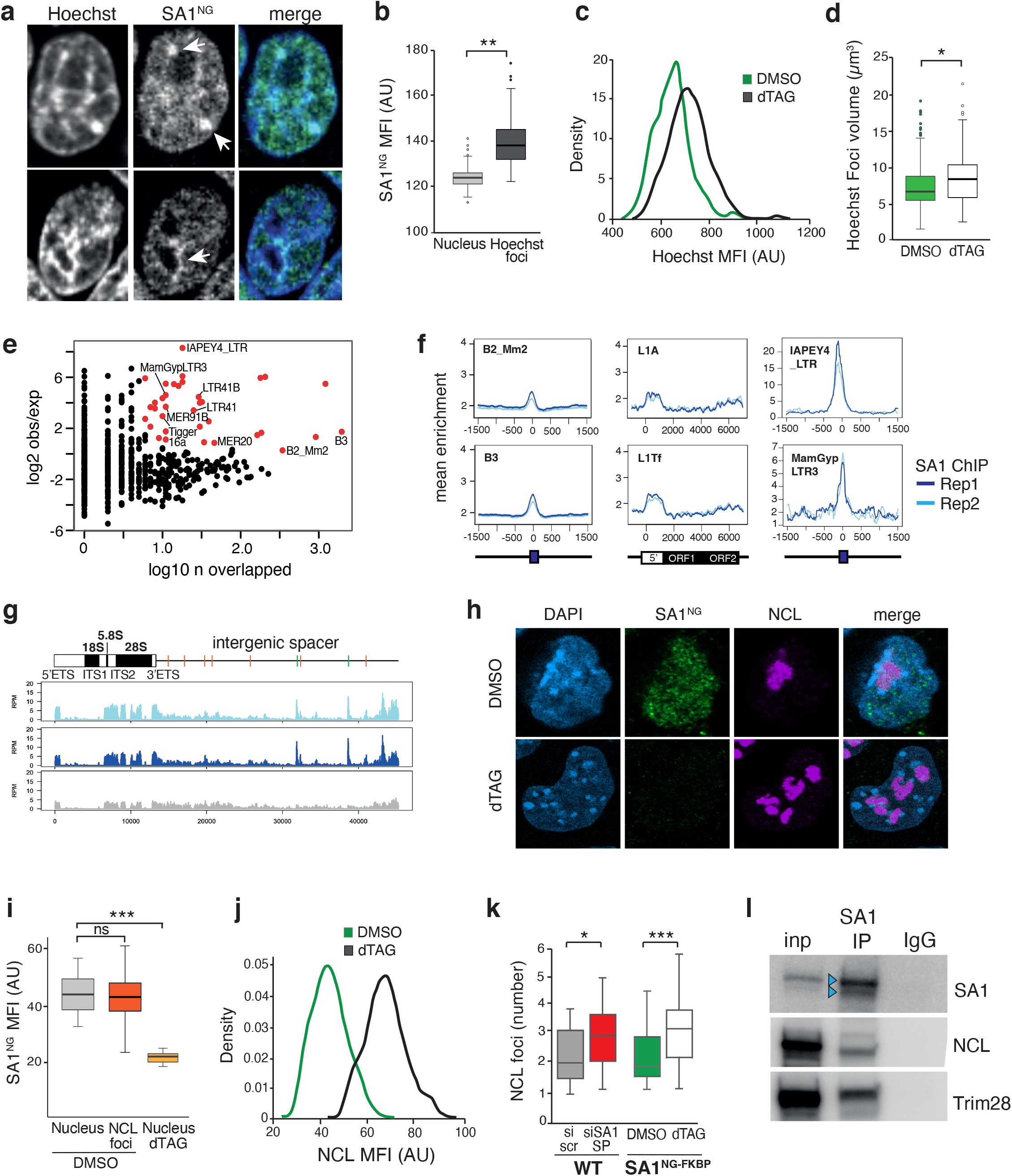
Stag1 is localised to and impacts both euchromatin and heterochromatin compartments. a) Live-cell Spinning Disk confocal images of two SA1^NG-FKBP^ mESCs counterstained with Hoechst. Arrows indicate notable regions of overlap of SA1 and Hoechst, including at Hoechst-dense foci and at the nucleolar periphery. *NB* Puncta within the nucleoplasm can also be observed. b) Imaris quantification of the MFI of SA1-NeonGreen within the nucleus (light grey) or Hoechst-dense foci (dark grey). Quantifications and statistical analysis were done as above. Data is from two independent experiments, n>50 cells/condition. AU, arbitrary units. c) Distribution of Hoechst MFI from SA1^NG-FKBP^ mESCs treated with DMSO (green) or dTAG (black). Data is from n>100 cells/condition. d) Imaris quantification of the volume of Hoechst foci in SA1^NG-FKBP^ mESC treated with DMSO (green) or dTAG (white). Quantifications and statistical analysis were done as above. Data is from two independent experiments, n>50 cells/condition. AU, arbitrary units. e) Number of copies of each repeat family that overlap a SA1 ChIP-seq peak and the enrichment of binding over random. Shown in red are the repeats which have significant enrichment, with a subset of these labelled. f) Profiles of the mean enrichment of SA1 ChIP-seq at select TE repeat families. Shown are full-length elements of the indicated SINE, LINE and LTR families. Two SA1 ChIP replicates are shown in blue. g) Top, cartoon of the consensus Mus musculus ribosomal DNA (rDNA) (GenBank: BK000964.3), showing the ribosomal genes and the intergenic spacer (IGS) region which contains several SINE elements (Red, B2_Mm2; Green, B3). Bottom, Stag1 ChIP replicates and INPUT as in f) above, aligned to this region. h) Representative confocal images of MFI of SA1-NeonGreen and Nucleolin (NCL) assessed by IF in SA1^NG-FKBP^ mESCs treated with DMSO or dTAG and counterstained with DAPI. i) Imaris quantification of the MFI of SA1-NeonGreen from h) within the nucleus or NCL foci in DMSO and dTAG conditions. Quantifications and statistical analysis were done as above. Data is from two independent experiments, n>50 cells/condition. AU, arbitrary units. j) Distribution of NCL MFI from SA1^NG-FKBP^ mESC treated with DMSO (green) or dTAG (black). Data is from n>100 cells/condition. k) Imaris quantification of the number of NCL foci in wildtype mESC treated with si scr (grey) or siSA1 SP siRNAs (red) and in the SA1^NG-FKBP^ mESC clone treated with DMSO (green) or dTAG (white). Quantifications and statistical analysis were done as above. Data is from two independent experiments, n>50 cells/condition. See also Figure S2. l) Chromatin immunoprecipitation of SA1 and IgG from wildtype mESCs and WB for SA1, NCL and Trim28. Blue arrows indicate multiple immunoreactive bands to SA1.

These observations prompted us to re-analyse STAG1 chromatin immunoprecipitation followed by sequencing (ChIP-seq) data in mESC ^26,55^. We calculated the proportion of STAG1 peaks that overlapped genes, repeats (within the Repeat Masker annotation), introns and intergenic regions not already represented (see Methods). Of the 18,600 STAG1 peaks identified, the majority (76%) are bound to genomic sites that are distinct from protein-coding genes including at repetitive elements and intergenic regions (Fig. S2c). Indeed, STAG1 binding was enriched at specific repeat families above random expectation (Fig. 2e). These included the DNA transposon and Retrotransposon classes, both known to form constitutive heterochromatin in differentiated cell types, are expressed in early development and involved in regulation of cell fate ^56,57^. Specifically, STAG1 was enriched at SINE B3 and B2-Mm2 elements (previously shown to be enriched at TAD borders ^58^); several LTR families, two of which have been previously shown to be associated with CTCF (LTR41, LTR55) ^59^ and at evolutionary young and active families of LINE1 elements (L1Tf, L1A) (Fig. 2e, f, S2e). We also found that several SINE B3 elements located within the intergenic spacer (IGS) of the consensus rDNA locus were bound by STAG1 (Fig. 2g). The binding of STAG1 at repeats may be dependent on CTCF since many of the bound repeats contained CTCF motifs (Fig. S2d).

RNA-seq of siSA1-treated mESC did not reveal dramatic changes in steady-state transcription of repetitive elements. However, qRT-PCR analysis using primers to ORF1 of Stag1-bound LINE1 and pre-rRNA revealed reduced expression compared to controls (Fig. S2f), suggesting a possible role for Stag1 in the control of repeat expression. Together with the microscopy results, the profile of STAG1 peaks suggests that the role of Stag1 in mESCs may extend beyond protein-coding gene regulation.

### STAG1 supports nucleolar structure

In mESCs, LINE1 transcripts have been shown to act as a nuclear RNA scaffold for the interaction with the nucleolar protein Nucleolin (NCL), a regulator of rRNA transcription, and the co-repressor TRIM28 (Kap1) ^60^. The complex promotes rRNA synthesis, nucleolar structure and self-renewal in mESC ^56^. Since depletion of Stag1 results in a loss of self-renewal and reduced rRNA expression and Stag1 was enriched at LINE1 and rDNA, we considered whether Stag1 was supporting pluripotency through nucleolar structure and function. We were not able to use spinning disk microscopy to assess the co-localization of Stag1 with nucleolar proteins in live cells. Instead, we used confocal imaging of SA1^NG_FKBP^ mESC stained with NCL. We observed a similar amount of SA1-NeonGreen (SA1^NG^) within the nucleolus compared to the nucleus of mESC (Fig. 2h, i). Notably, upon dTAG-treatment of SA1^NG_FKBP^ mESC, there was a significant increase in NCL signal intensity (Fig. 2j) as well as increased numbers of nucleolar foci in both dTAG-treated SA1^NG_FKBP^ and in siSA1 KD mESCs (Fig. 2k, S2g, h), reminiscent of changes observed during mESC differentiation ^61^. Further, STAG1 immunoprecipitation followed by WB in mESC revealed an interaction with both NCL and Trim28 (Fig. 2l), suggesting a direct effect of Stag1 on nucleolar structure and rRNA expression.

### Stag1 expression is highly regulated in mESCs

We consistently observed several immunoreactive bands on Stag1 WB (Fig. 2l, arrows), which were enriched in mESC (Fig. 1b). In order to gain a full perspective on how Stag1 may be contributing to nucleolar structure and pluripotency, we first investigated whether *STAG1* may be regulated at the level of transcription in mESCs. Several lines of evidence suggested that this may be the case. First, STAG1 levels are higher in 2i-grown compared to FCS-grown mESCs, a culture condition that supports a mix of naïve and primed cells (Fig. S1b, d) and second, primers positioned along the length of STAG1 amplify mRNAs that respond differently to differentiation (Fig. 1a). Thus, we employed a series of approaches to comprehensively characterize Stag1 mRNAs. First, we used RACE (Rapid Amplification of cDNA Ends) to characterize the starts and ends of Stag1 mRNAs directly from mESCs. 5’ RACE uncovered four novel alternative transcription start sites (TSS) in mESCs; ∼50kb upstream of the canonical Stag1 TSS (referred to as ‘SATS’, and previously identified in ^62^) (Fig. 3a, d, S3a); between canonical exon 1 and exon 2 (referred to as alternative exon 1 or altex1) (Fig. 3a, d, S3d); and at exons 6 and 7 (Fig. 3a, d, S3a). Interestingly, the TSS located at exon 7 (e7) was preceded by a sequence located *in trans* to the STAG1 gene, carrying simple repeats and transcription factor binding sites (Fig. 3b). While the frequency of this alternative TSS was significantly lower than the other TSSs, it was identified in multiple RACE replicates, indicating that it may be present in a subset of the mESC population. We also discovered widespread alternative splicing in the 5’ region of Stag1, with particularly frequent skipping of exons 2 and 3 (e2/3Δ) and exon 5 (e5Δ) (Fig. 3d, S3a, f). Using 3’ RACE, we detected an early termination site in intron 25 and inclusion of an alternative exon 22 introducing an early STOP codon, as well as several 3’UTRs (Fig. 3 c, d, S3c).

**Figure 3.**
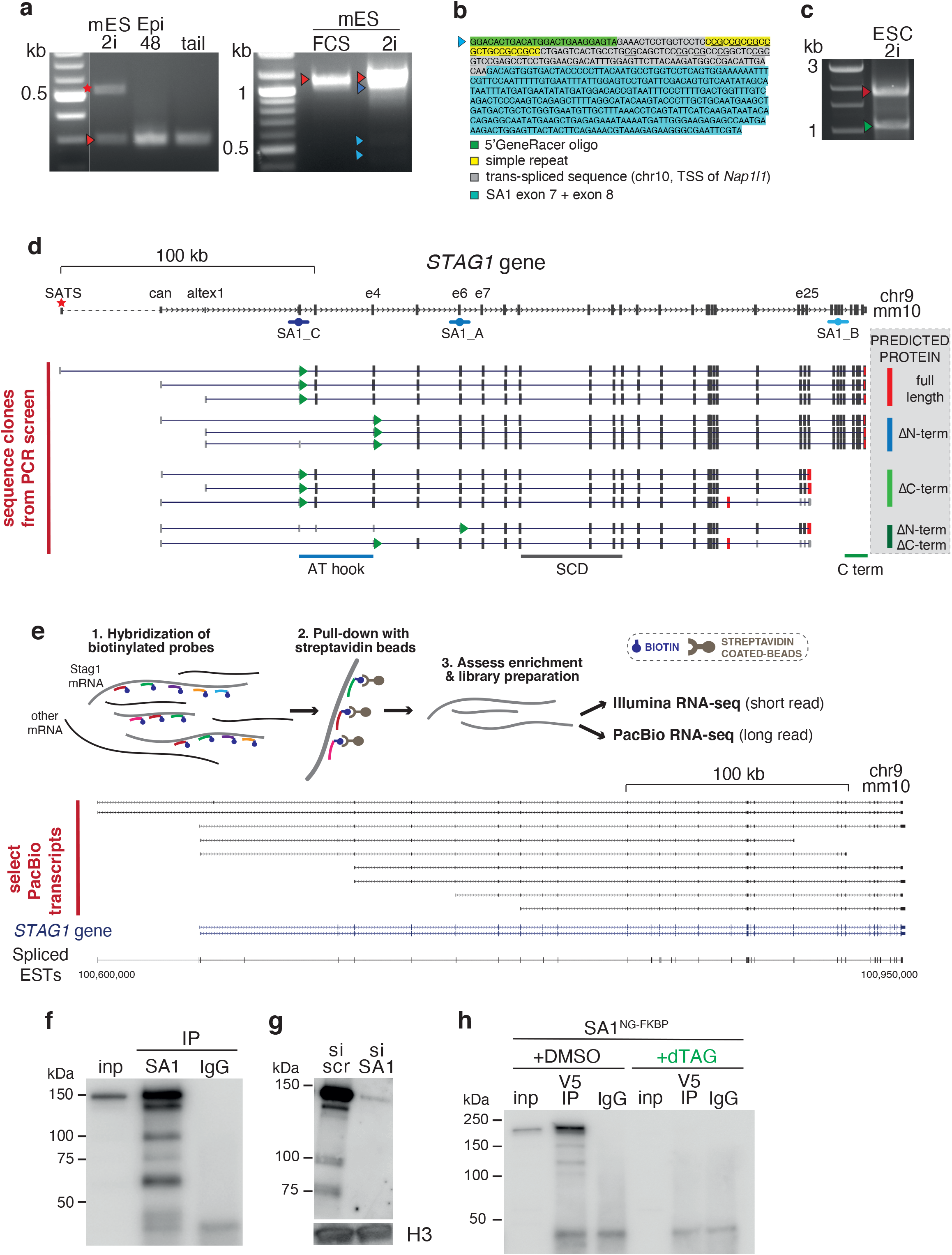
Stag1 undergoes widespread transcriptional regulation in mESCs. a) 5’ Rapid Amplification of cDNA ends (RACE) for SA1 in naïve mESC and EpiLCs. Left gel; red star indicates SATS TSS and red arrow indicates canonical (can) TSS. Right gel; red arrow indicates full length Stag1 with both SATS and can TSSs; dark blue arrow indicates alternatively spliced variants arising from skipping of exons in the 5’ region; light blue arrows indicate the TSSs at exon 6 (e6) and exon 7 (e7). Arrows indicate bands which were cloned and sequenced. See also Figure S3. b) The 5’ RACE fragment that identified a new TSS at exon 7 spliced directly to a sequence in trans carrying regulatory elements. c) 3’ RACE for SA1 in naïve mESCs. Red arrow indicates canonical full-length end; green arrow indicates end in i25. Arrows indicate bands which were cloned and sequenced. See also Figure S3. d) Top, schematic of the *STAG1* gene annotation in mm10. The identified TSS and TTSs from RACE are indicated. Bottom, aligned sequence clones from the PCR mini-screen and their predicted impact on the SA1 protein (grey box, right). Green arrows and red bars within the transcripts indicate start of the coding sequence and the TTS respectively. Shown also are the regions which code for the AT hook and the stromalin conserved domain (SCD). e) Schematic of the PacBio sequencing methodology (see methods for full description). Select transcripts sequenced on the PacBio platform, including many isoforms already discovered using RACE and PCR cloning methods above. See also Figure S3. f) WB analysis of endogenous, chromatin-bound SA1 protein isoforms from mESCs and g) upon treatment with si scr and siSA1. H3 serves as a loading control. h) Chromatin immunoprecipitation for the v5 tag in SA1^NG-FKBP^ mESCs treated with DMSO or dTAG to degrade SA1. *NB*. SA1 bands run 42kDa higher due to the addition of the tag.

Next, PCR- and Sanger sequencing-based clonal screening confirmed that the newly discovered 5’ and 3’ ends represent true Stag1 transcript ends, validated the existence of the e2/3Δ and e5Δ isoforms, confirmed their enrichment in naïve mESCs compared to differentiated mouse embryonic fibroblasts (MEFs) and uncovered an isoform lacking exon 31 which encodes a basic domain embedded in the otherwise acidic C-terminal region of Stag1 (e31Δ) (Fig. S3d). To determine the complete sequences of the Stag1 transcript isoforms and to use a non-PCR-based approach, we performed long-read PacBio Iso-seq from 2i mESC RNA (Fig. 3e). This confirmed the diversity of the Stag1 5’ and 3’UTRs, the e31Δ isoform, multiple TSSs including SATS, and early termination events, including in i22 and i25 (Fig. 3e, S3e). Importantly, these transcripts all had polyA tails, in support of their protein-coding potential. Finally, we validated and quantified the newly discovered splicing events by calculating the frequency (percentage spliced in (PSI)) of exon splicing in our RNA-seq as well as in published data using the VAST-tools method ^63^. This confirmed the presence of Stag1 splicing events in other mESC datasets and supported that several of these were specifically enriched in mESC (Fig. S3f, Table S1).

Interestingly, visual inspection of the genome topology around the *Stag1* locus in our 2i mESC and neural stem cell (NSC) Hi-C data ^64^ revealed that the *STAG1* gene undergoes significant 3D reorganization as cells differentiate (Fig. S4). For example, the *STAG1* TAD switches from the active to the repressive compartment during differentiation, in line with the decrease in Stag1 levels during differentiation. Furthermore, UMI-4C revealed changes to sub-TAD architecture corresponding to the newly discovered mESC-enriched Stag1 TSSs and TTSs described above, suggesting that 3D chromatin topology may play a role in facilitating the transcriptional diversity of *STAG1* (Fig. S4). Together, our results point to a previously unappreciated diversity of endogenous Stag1 transcripts in mESCs, prompting us to investigate the importance of these for pluripotency and the nucleolus.

### Multiple Stag1 protein isoforms are expressed in mESCs

Stag1 transcript diversity was intriguing because many of the events were either specific to mESC or enriched compared to MEFs and NSCs (Fig. S3d, f). Furthermore, the transcript variants were predicted to produce STAG1 protein isoforms with distinct structural features and molecular weights (Fig. 3d, S3g). For example, the truncation of the N-terminus (e2/3Δ, e5Δ, e6 TSS and e7 TSS), and thus loss of the AT-hook (amino acid 3-58), could impact STAG1 association with nucleic acids. Meanwhile, C-terminal truncated Stag1 isoforms (altex22, i25 end, e31Δ) could affect STAG1-cohesin interactions. It is noteworthy that the evolutionarily conserved Stag-domain (‘SCD’, AA 296-381) ^30^, shown to play a role in CTCF interaction ^29^, would be retained in all the isoforms identified here.

Immunoprecipitation (IP) of endogenous STAG1 followed by WB revealed multiple bands corresponding to the predicted molecular weights for several protein isoforms and identified by mass spectrometry to contain Stag1 peptides (Fig. 3e, S3g, Table S2). Similarly, multiple bands of expected sizes were reduced between naïve and primed cells (Fig. S3h) and sensitive to Stag1 KD, alongside the canonical, full-length isoform (Fig. 3f). Treatment of SA1^NG_FKBP^ mESCs with dTAG followed by WB of chromatin-associated proteins with an antibody to the v5 tag further confirmed the sensitivity of the isoforms to dTAG-mediated degradation (Fig. 3g). Thus, complex transcriptional regulation in mESCs gives rise to multiple Stag1 transcripts and protein isoforms with distinct regulatory regions and coding potential. Our discovery of such naturally occurring isoforms offers a unique opportunity to define the functions of the divergent N- and C-terminal ends of Stag1 in the context of the pluripotent state.

To study the functional consequences of the Stag1 isoforms on pluripotency and nucleolar structure, we took advantage of our detailed understanding of Stag1 transcript diversity to design custom siRNAs to selectively target, or retain specific isoforms (Fig. 4a). Alongside the siRNAs used in Figure 1 (SmartPool, SP), we designed siRNAs to specifically target the SATS 5’UTR (esiSATS), the 5’ end (siSA1-5p) or the 3’ end (siSA1-3p) of Stag1 mRNA (see Methods). We anticipated that the KD panels would not completely abolish all Stag1 transcript variants, but rather change the relative proportions, in effect experimentally skewing the levels of the N- and C-terminal ends of Stag1 in cells. 3p siRNAs were predicted to downregulate full-length and N-term truncated isoforms and retain C-term truncated isoforms, while 5p siRNAs would specifically retain N-term truncated isoforms.

**Figure 4.**
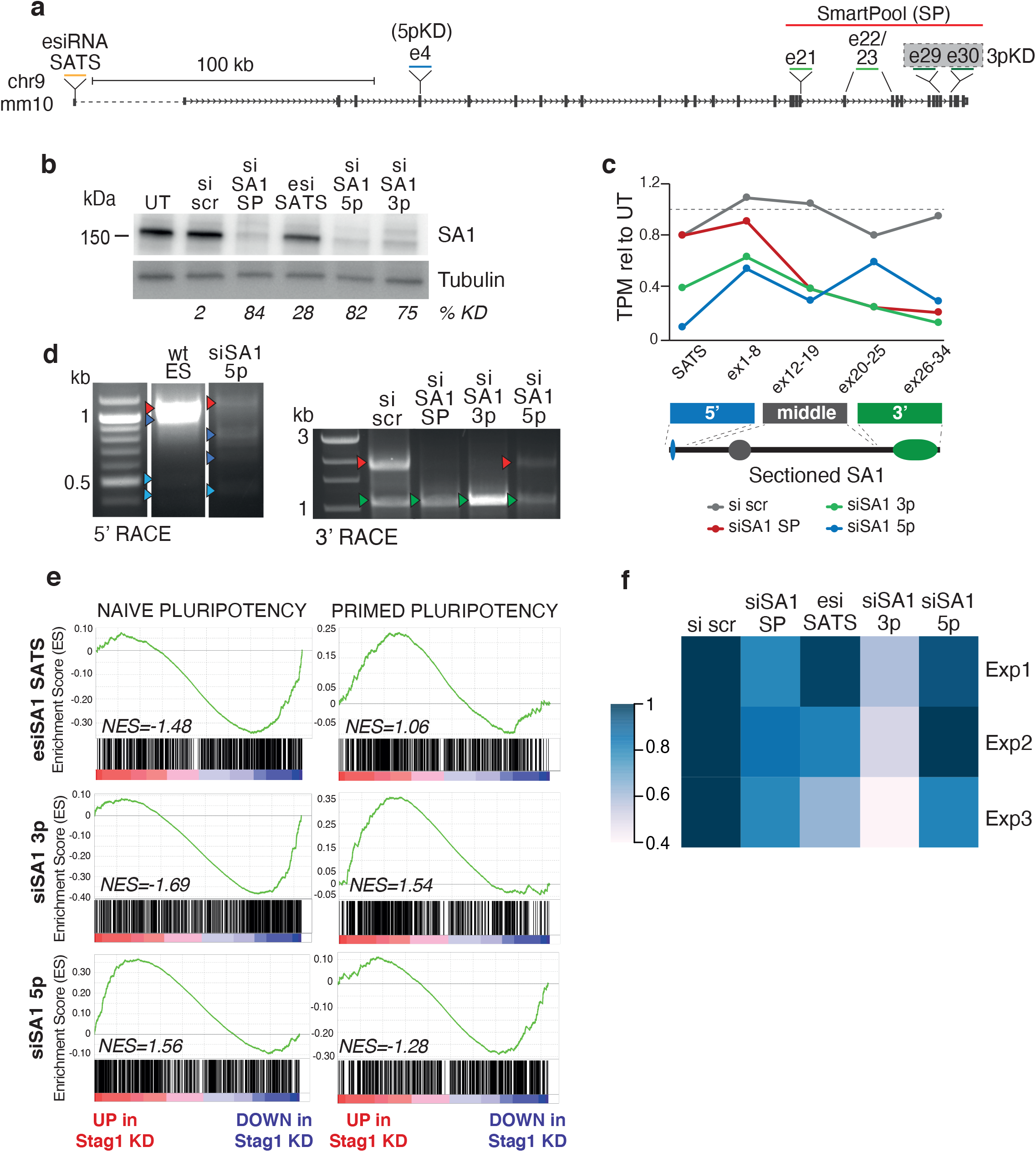
Fluctuations in the levels of the Stag1 isoforms skews cell fates. a) Schematic of the siRNA pools used in this study. esiRNA SATS represents ‘enzymatically-prepared’ siRNAs (see Methods). b) WB analysis of SA1 levels in mESC WCL after no treatment (UT), or upon si scr, si SA1 SP, si SA1 3p, si SA1 5p or esi SATS treatment. Tubulin serves as a loading control. The percentage of knockdown (KD) of SA1 signal normalised to Tubulin is shown. c) RNA-seq reads (TPM, transcripts per million) aligning to sectioned Stag1 in datasets from the various siRNA pools, shown as relative to untreated mESC RNA-seq. N-terminal reads include SATS and exons 1-8, Mid reads include exons 12-19 and C-terminal reads include exons 20-25 and exons 26-34. *NB*. the change in read proportions in the different KD treatments. d) Left gel, 5’ and Right gel, 3’ RACE for SA1 in mESC treated with the indicated siRNAs. Arrows indicate bands which were cloned and sequenced and colour-coded as before. e) Enrichment score (ES) plots from GSEA using the naïve and primed gene sets as in Fig. 1e and RNA-seq data from the indicated siRNA treated mESC samples. f) Area occupied by AP+ colonies in mESC treated with the siRNA panel from three independent biological replicates. n>50 colonies/condition were counted.

siRNAs to the 5p and 3p ends of Stag1 reduce full-length Stag1 mRNA and protein with similar efficiency to SP KDs. esiSATS reduces Stag1 by ∼30-50%, indicating that the SATS TSS functions to enhance expression of Stag1 in naïve mESC (Fig. 4b, S5a). We confirmed that Stag1 isoform proportions were altered upon siRNA treatment using RNA-seq, RACE and immunoprecipitation. RNA-seq reads aligning to Stag1 in the different siRNA treatments were quantified to represent the residual N-terminal, middle and C-terminal read proportions (Fig. 4c). Residual reads in the SP and 3p KDs aligned predominantly to the N-terminus and were depleted from the C-terminus. While the 5p KD had the least read retention in the N-terminus (Fig. 4c). In parallel, we performed RACE to validate changes to the proportions of Stag1 isoforms. 5’ RACE performed in mESC treated with 5p siRNA revealed downregulation of full-length Stag1 transcript while several N-terminal truncated isoforms were upregulated compared to untreated cells (Fig. 4d, left panel, blue arrows). Similarly, transcripts terminating at the canonical 3’ end of Stag1 are strongly reduced in the SP and 3p siRNA KD samples and to a lesser extent in the 5p KD (Fig. 4d, red arrows), supporting the expectation that residual transcripts in the 5p KD have C-terminal ends. Meanwhile, the transcript terminating in i25 is substantially enriched upon 3p KD (Fig. 4d, right panel, green arrows). Thus, the siRNA panel developed here provide us with a powerful tool to modulate the proportion of the naturally occurring Stag1 isoforms in mESCs and study their potential roles in pluripotency.

### A specific role for the Stag1 C-terminus in the maintenance of naïve pluripotency transcriptome

We first quantified the effect of the Stag1 siRNA KDs on pluripotency gene expression. qRT-PCR for Nanog expression and WB for Nanog protein levels revealed that the 3p KD had a similar effect on Nanog to SP, with significant downregulation, while surprisingly, the 5p KD did not reduce Nanog (Fig. S5b). We prepared biological replicate RNA-seq libraries from the Stag1 3p, 5p and SATS siRNA KDs. We used GSEA as before to probe for signatures of naïve or primed pluripotency. In support of our previous results, reducing Stag1 levels by targeting the mESC-specific SATS promoter leads to downregulation of the naïve pluripotency gene signature and upregulation of the primed signature (Fig. 4e, S5c), reminiscent of the phenotype from SP KD (Fig. 1e, f). We again observed a differential effect of the 3p and 5p KDs on naïve and primed pluripotency signatures. A similar but more prominent loss of the naïve signature was observed in 3p KD RNA-seq compared to SATS and SP, while surprisingly, in 5p KD cells the naïve signature was unaffected compared to si scr controls (Fig. 4e).

The distinct gene expression profiles of the 3p and 5p KDs were reflected in differences in self-renewal. Cells treated with 3p siRNAs exhibited a significant loss of self-renewal potential, consistent with the loss of the naïve pluripotency signature, with only 20% of colonies exhibiting AP-staining compared to 30% of colonies in the SP KDs (Fig. S5d), and an average reduction of the area occupied by AP+ colonies of 50% compared to si scr controls (Fig. 4f). This was not evident in the 5p KD, where the effect on self-renewal was more similar to si scr controls (Fig. 4f). Interestingly, unlike siRNA to Stag1, esiSATS results in a variable effect on self-renewal (ranging from between 5-35% reduction in AP+ area) (Fig. 4f), likely because the SATS TSS is expressed in the most naïve cells of the population, the frequency of which varies significantly between FCS populations. Our results further confirm the importance of Stag1 in self-renewal and point to a specific role for the C-terminal of Stag1 in maintaining a naïve pluripotency gene expression programme.

### The N-terminus of Stag1 supports nucleolar structure and function

The different effect on naïve pluripotency between the 3p and 5p KDs was surprising. We therefore sought to re-examine the effect of our siRNA panel on the Stag1 bound repeats LINE1 and rDNA (Fig. 2f, g). As we had not observed a significant difference on steady state levels of repeats from our RNA-seq experiments, we instead purified nascent RNA from mESCs treated with siRNAs. Both the KD and the nascent RNA pull-downs were successful as revealed by qRT-PCR to Stag1 (Fig. 5a, b). Consistent with our previous results, total Nanog RNA levels were significantly reduced in siSA1 SP and 3p KD but not in 5p KD. Interestingly, this trend was not observed in nascent levels of Nanog RNA where the 3p KD does not have a significant effect, suggesting that the C-terminus may be required for the stability of Nanog mRNA instead of its transcription *per se* (Fig. 5a, b). Upon Stag1 SP KD, both steady state and nascent levels of LINE1 RNA were modestly decreased (also Fig. S2f). While the 3p KD had a 20% reduction in LINE1 RNA expression, this was not maintained at steady state levels. However, both nascent and total levels of LINE1 RNA were significantly reduced by 40-50% of controls in 5p KD mESCs. These results were also observed for pre-rRNA, with only the SP and 5p KD having significant effects on expression. Thus, the N-terminus of Stag1 plays a distinct role in LINE1 and rDNA expression (Fig. 5a, b).

**Figure 5.**
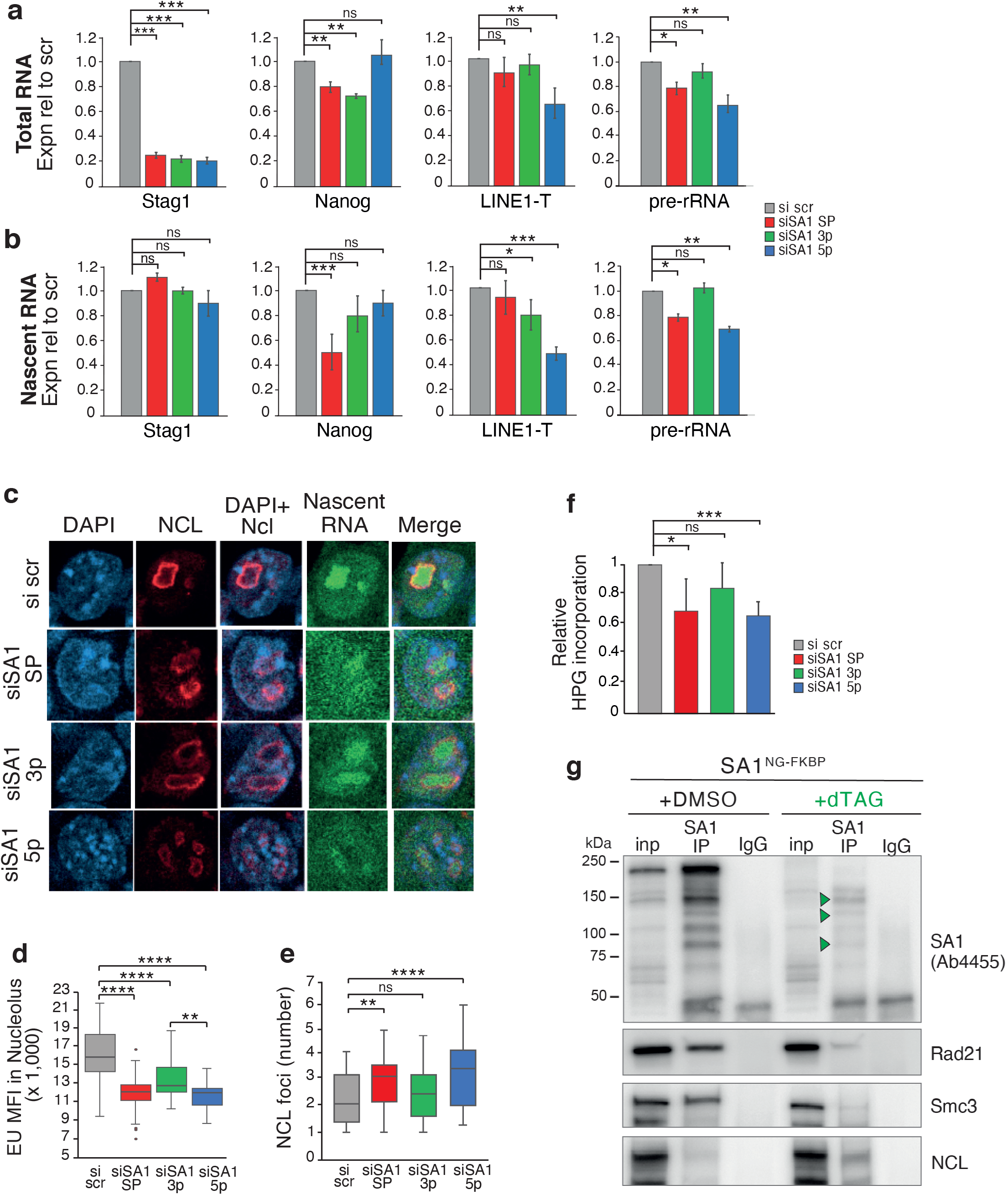
The N- and C-terminal ends of Stag1 regulate expression in different genomic compartments. Relative expression of Stag1, Nanog, LINE1-T and pre-rRNA by qRT-PCR in mESC after treatment with the siRNA panel. Shown are a) total and b) nascent RNA levels. Data is represented as mean ± SEM and statistical analysis as before. Data is from three independent experiments. c) Representative confocal images of IF to NCL and nascent RNA in siRNA-treated mESC labelled with EU-488. Nuclei were counterstained with DAPI. d) Imaris quantification of the MFI of nascent RNA (EU) within the nucleoli from (c), as defined by a mask made to the NCL IF signal. Quantifications and statistical analysis were done as above. Data is from two independent biological replicates. n>50/condition, except for siSA1 5p where n>35. e) Imaris quantification of the number of NCL foci in siRNA-treated mESCs. Quantifications and statistical analysis were done as above. Data is from two independent experiments, n>50 cells/condition. f) Analysis of global levels of nascent translation by measuring HPG incorporation using Flow cytometry and analysed using FloJo software. Shown is the quantification of the change in EU incorporation relative to si scr treated cells. Data are from four biological replicates. g) Chromatin immunoprecipitation using an N-terminal Stag1 antibody in SA1^NG-FKBP^ mESC treated with DMSO or dTAG. Green arrow indicates residual C-terminal truncated Stag1 isoforms. Shown also are WB for the core cohesin subunits Rad21 and Smc3 and NCL.

Given the effects on LINE1 and rRNA, we also assessed nucleolar structure and function using our siRNA panel. mESC were pulsed with 5-ethynyl uridine (EU) which becomes actively incorporated into nascent RNA and enables detection of newly synthesized RNA. Samples for IF were co-stained with an antibody to NCL to simultaneously quantify nucleoli number and changes in nascent RNA transcription. Cells treated with scrambled siRNA showed a distinct nucleolar structure and the EU signal could be seen throughout the nucleus, with a strong enrichment within the nucleolus as expected from rRNA expression (Fig. 5c). While a significant reduction in nascent RNA signal was observed in all KD conditions compared to scrambled controls (Fig. S5e), by IF, we observed a distinct effect on nascent RNA levels within the nucleolus in the 5p KD. While the medians between the three siSA1 KDs were not dramatically different, the effect of the 5p KD on nucleolar RNA signal distribution was significantly different from the 3p KD (Fig. 5d). This result was consistent with the qRT-PCR analysis of nascent pre-rRNA levels (Fig. 5b) and with the significant effect on NCL foci number in 5p KD mESCs (Fig. 5e). Consequently, we also observed changes to global translation by assessing the incorporation of L-homopropargylglycine (HPG), an amino acid analogue of methionine into mESC using FACS analysis. HPG incorporation was significantly reduced in SP and 5p siRNA treated mESCs compared to scrambled control (32% and 35% of si Scr) (Fig. 5f, S5f). We did observe a modest effect on global nascent translation in 3p KD treated cells (16% of si scr), although this was not significantly different from scrambled control. Our results reveal distinct roles for the N- and C-termini of Stag1 in nucleolar structure and function and pluripotency gene expression, respectively.

The effects observed on rRNA levels and nucleolar function were not associated with changes to expression of ribosome subunit expression (Fig S5g). Thus, we considered whether the regulation of LINE1 expression by the N-terminus of Stag1 influenced nucleolar structure via the NCL/Trim28 complex (Fig. 2l). To investigate this, we took advantage of our Stag1^NG_FKBP^ mESCs. dTAG treatment can only degrade isoforms containing the FKBP tag inserted into the canonical C-terminal end. Thus Stag1^NG_FKBP^ mESCs treated with dTAG should enrich for SA1^ΔC^ isoforms which contain an N-terminus. Indeed, immunoprecipitation of STAG1 using an antibody which recognizes an N-terminal epitope reveals the presence of several N-terminal-enriched SA1ΔC isoforms (Fig. 5g, green arrows). WB of this IP material revealed a reduction in the ability of SA1ΔC to interact with the cohesin subunits Rad21 and Smc3, despite similar levels in the input of dTAG treated cells. Meanwhile, the interaction with NCL was increased in same lysate (Fig. 5g). Taken together, our results are supportive of the different ends of Stag1 interacting with different protein partners to co-ordinately regulate pluripotency.

### The N-terminus of Stag1 suppresses the totipotent state

In addition to promoting rRNA synthesis and self-renewal in mESC, the LINE-1/NCL/Trim28 complex represses a transcriptional program specific to totipotent cells in the two-cell (2C) stage of development, termed two-cell-like (2C-LC) ^56^. The phenotypes of the 5p KD, namely reduced rRNA and LINE-1 expression, reduced translation and aberrant nucleolar function, pointed towards possible conversion of cells into a 2C-LC state. We therefore tested whether Stag1, and specifically the N-terminal end, play a role in totipotency.

We first investigated whether 2C-L cells which naturally arise within mESC populations express Stag1NΔ isoforms. To formally address this, we obtained mESCs expressing a Dox-inducible *Dux-HA*-expression construct together with a MERVL-linked GFP reporter ^65^. Dux is a 2C-specific transcription factor which binds to MERVL elements to activate expression (Hendrickson et al., 2017). We induced *DuxHA-*expression in the MERVL-GFP mESC and performed 5’ RACE as before on sorted GFP+ (2C-L) and GFP-cells (Fig. 6a). We enriched several of the previously identified N-term truncated Stag1 transcripts in the GFP+ population including e2/3Δ and e5Δ isoforms (Fig. 6a, blue arrows). Importantly, we also identified a transcript starting at e7, similar to the one previously found in 5p KD mESC (Fig. 6b, 3a, b). Remarkably however, the sequence preceding the TSS in e7 in *Dux*-induced cells was an MT2-MERVL element, creating a chimeric, LTR-driven Stag1 transcript, reminiscent of other LTR-transcripts specifically expressed in the 2C-L state.

**Figure 6.**
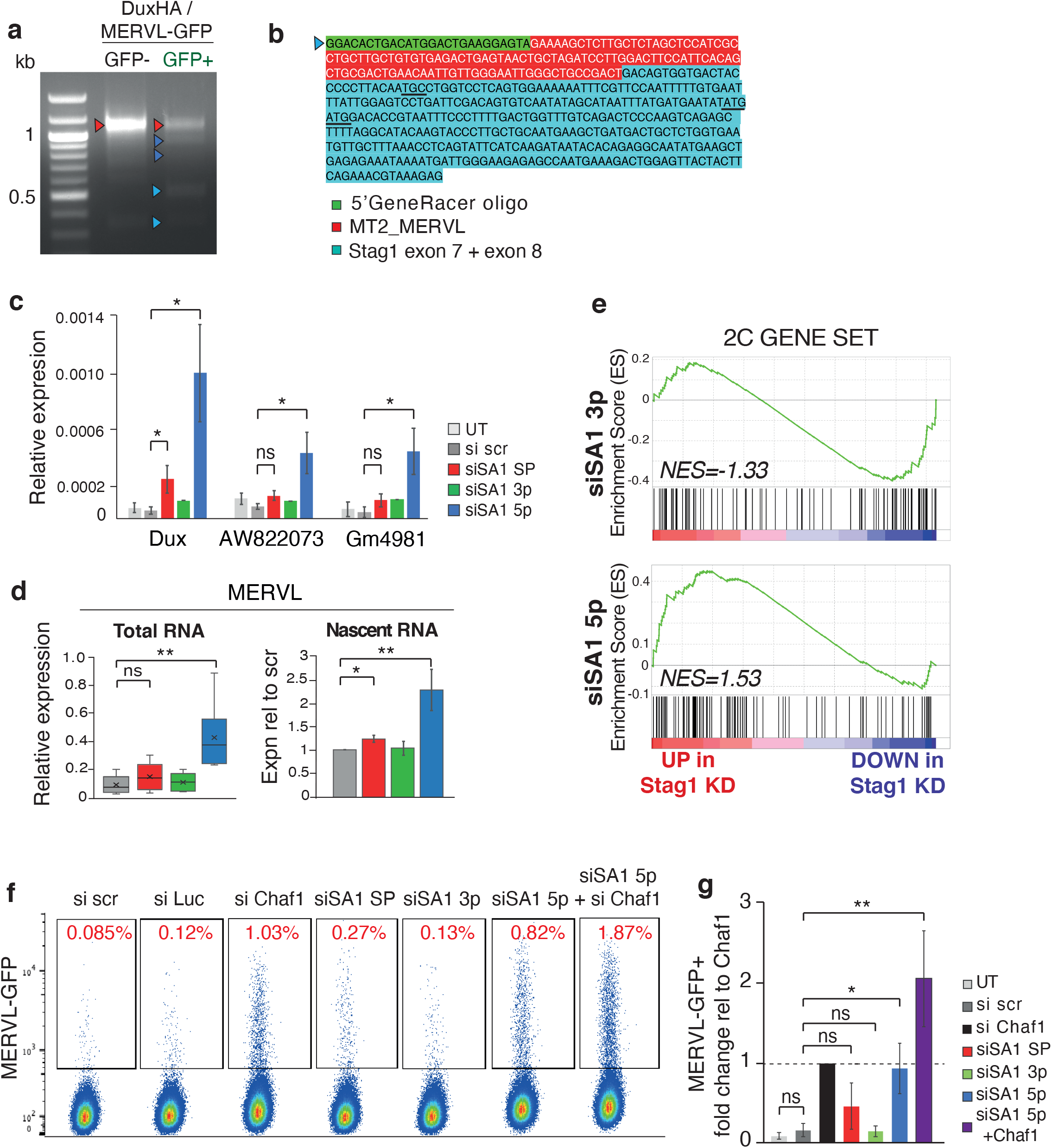
Stag1 N-terminus protects against conversion of ESCs to totipotency. a) 5’ RACE for Stag1 in Dux-HA MERVL-GFP mESCs with and without sorting for GFP+ cells. Arrows indicate bands which were cloned and sequenced and colour-coded as previously described. b) Sequence of the 5’RACE product identifying a novel Stag1 TSS from (a) with direct splicing of exon7 to an MT2_MERVL element. c) Relative expression of several 2C-LC markers in total RNA by qRT-PCR in mESC after treatment with the siRNA panel. Data is represented as mean ± SEM and statistical analysis as before. Data is from six independent experiments. d) Relative expression of MERVL repeat element by qRT-PCR in mESC after treatment with the siRNA panel. Shown are total (left) and nascent RNA (right) levels. Quantifications and statistical analysis as before. Data is from five biological replicates. *NB*, nascent RNA levels are shown relative to si scr control. e) Enrichment score (ES) plots from GSEA using a published 2C-L gene set and RNA-seq data from the 3p and 5p siRNA treated mESC samples used in Figure 4. f) Representative FACS analysis of the proportion of mESCs expressing a MERVL-GFP reporter in the different siRNA treated cells and including siRNA to Chaf1 as a positive control. Percentage of MERVL-GFP+ cells based on Flo-Jo analysis is shown in red. g) Proportion of MERVL-GFP+ cells in the different siRNA conditions relative to the siChaf1 positive control. Data is represented as mean ± SEM and statistical analysis as before and is from four independent experiments.

2C-LCs are a rare subpopulation which spontaneously arise in mESC cell cultures and exhibit unique molecular and transcriptional features ^43,66,67^. Given that 2C-LCs expressed several N-term truncated Stag1 isoforms, we investigated whether these in turn supported the maintenance or emergence of that state. We treated mESCs with the panel of siRNAs and used RT-qPCR to test expression of candidate genes. We found that Dux, and consequently MERVL and other markers of the totipotent 2C-L state, Gm6763, AW822073 and Gm4981 are strongly upregulated by 5p KD (Fig. 6c, d, S6a). Notably, all 2C-L genes analysed remained unchanged in 3p KD conditions with a modest upregulation in SP KD. Further, GSEA using a published 2C gene set ^56^ revealed a specific enrichment among the upregulated genes in 5p KDs that was not observed in 3p KDs (Fig. 6e, S6b), consistent with the different ends of Stag1 targeting different RNA pools.

To functionally validate the expression results, we returned to the Dox-inducible *Dux-HA*, MERVL-GFP mESCs ^65^ and used flow cytometry to directly measure the number of GFP-positive cells in our different Stag1 KD conditions (Fig. 6f, g). Chaf1 is a chromatin accessibility factor previously shown to support conversion of mESC towards totipotency ^43^. In support of the upregulation of the 2C-LC gene set in 5p KD mESCs, we observed an 8-9-fold increase in the proportion of GFP-positive cells in 5p KD conditions compared to scramble treated controls, similar to the published effect of Chaf1 KD (Fig. 6f, g). There was a modest, but insignificant increase in GFP+ cells upon SP KD and no effect upon 3p KD. mESC treated with both Chaf1 and 5p siRNAs had an additive effect on the proportion of GFP-positive cells, suggesting that the two proteins function in complementary pathways for conversion towards totipotency. Thus, 2C-LCs express N-term truncated Stag1 isoforms which in turn support the maintenance or emergence of that state through rRNA repression and nucleolar changes. Together our results reveal a new and specific role for the N-terminus of STAG1 in the regulation of the totipotent state.

## DISCUSSION

Most studies of cohesin function focus on the core trimer, despite the fact that it is the regulatory Stag subunit that are pan-cancer targets ^3^ and have clear roles in cell identity control ^2^. How these proteins contribute to cohesin’s functions, why cells have diversified them so extensively and how their mutations lead so often to disease are poorly understood. Here we reveal a novel role for Stag1, and in particular its unique N-terminal end, in regulating nucleolar integrity and 2C repression to maintain mESC identity. It has been known for a long time that several Stag paralogs exist in mammalian cells and that they have non-reciprocal functions with respect to chromosome structure and cohesion. By dissecting the diversity of naturally occurring Stag1 isoforms in mESCs, we have shed new light not only on the unique divergent ends of the Stag paralogs but also the critical role that their levels play in cell fate control. Our results highlight the importance of careful understanding of chromatin regulators in cell-specific contexts.

Stag1 knockout (Stag1^Δ/Δ^) ESCs give rise to mice which survive to E13.5 ^33,68^. At first this observation seems at odds with our report that Stag1 is required for pluripotency. However, our observations may in fact explain why the Stag1^Δ/Δ^ mouse model does not exhibit early embryonic lethality. In this model, only the 5’ region of Stag1 was targeted, meaning that the Stag1 isoforms lacking the N-terminus may still be retained in the targeted ESCs. This is consistent with our results showing that 5p KD cells have not lost their ability to self-renew nor is their pluripotency gene signature affected. It further suggests that changes to the nucleolus may exist in these cells.

The nucleolus is held together by liquid–liquid phase separation (PS), which is driven by the association of rDNA with nucleolar proteins and is dependent on continual rRNA synthesis ^37,38^. However, in one-to two-cell embryos, nucleoli lack distinct compartments, exhibit low rRNA synthesis and low translation ^69^. Similarly, changes to rRNA synthesis or nucleolar PS are sufficient to convert ESCs towards the 2C-LC state, either through Dux dissociation from the nucleolar periphery and consequently its de-repression ^44^ or p53-mediated nucleolar stress ^47^. Other proteins including the NCL/TRIM28 complex ^56^ and nucleolar LIN28 ^48^ have been shown to contribute to nucleolar integrity and repress DUX expression. In this context, our results position Stag1, and specifically its N-terminal end, as a novel regulator of the 2C-ESC transition through the control of nucleolar integrity. Stag1 is localised to the nucleolar periphery and interacts with the nucleolar proteins NCL/TRIM28 as well as being bound to and supporting rDNA and LINE-1 element expression. Our results suggest that the N-terminus of Stag1 plays a specific role in repressing conversion to 2C state. Stag1 may contribute to nucleolar structure and function via both the regulation of rRNA expression as well as by supporting nucleolar PS through interactions with nucleolar regulators. In this context, modulating the availability of the N- or C-terminus of Stag1 may be a way in which ESCs impact nucleolar structure and function and thus cell identity. Our results also point to the different ends of Stag1 interacting with different protein partners since mESCs retaining the C-terminus of Stag1 do not exhibit changes to the nucleolus and do not convert into 2C-LCs. This is also supported by the different gene expression programmes affected in the KDs that select for N-termΔ or C-termΔ isoforms. It may in fact be quite important for ESCs to express a diversity of alternative Stag1 isoforms to support plasticity of nucleolar structure and a range of cell fate options from totipotency to primed pluripotency.

Finally, Stag genes are commonly mutated in cancers ^3^. Our results point to misregulation of Stag proteins as leading to epigenetic misregulation, not necessarily only through changes to TADs and protein coding genes, but support a role for cell fate changes as a result of hierarchical changes to chromatin organization, nucleolar structure and function and repeat misregulation. Careful analysis of Stag2-mutant cancers should shed light on these and deliver new insights into cancers that harbour these mutations.

## Acknowledgments

This work would not be possible without the support of a Senior Research Fellowship from the Wellcome Trust awarded to S.H. (106985/Z/15/Z). We would like to thank Sally Lowell and Mattias Malaguti for advice throughout the project. We are grateful to the members of the Hadjur lab for critical discussions and reading of the manuscript. We thank M. Irima for help with VAST-tools pipelines; B. Cairns for the inducible Dux-HA, MERVL-GFP ESCs; W.Reik for a second MERVL-reporter ESC line; H. Rowe for advice on 2C-LC cells and repeat analysis. Thank you to Y. Guo and J. Manji in the Cancer Institute CRUK Centre FACS and Imaging core facilities for their invaluable assistance.

## Author Contributions

D.P. and S.H. conceived the project. D.P. designed and performed all the experiments on ESCs with assistance from S.W. S.W. performed all protein analysis, generated the SA1-NG-FKBP ESC line, performed the Spinning Disk microscopy and helped with the siRNA knockdown experiments. W.V. performed all bioinformatic analyses with the exception of the Stag1 enrichments at repeat elements, which was done by M.B. P.D. and S.P. provided advice on CRISPR targeting. D.P. and S.H. formatted all figures and wrote the manuscript with input from all authors.

## METHODS

### Embryonic stem cell culture and siRNA-mediated knockdown

Male mouse E14 embryonic stem cells (mESC) were cultured in serum (FCS) or naïve (2i) conditions. Serum-cultured cells were grown on 0.1% gelatin-coated plates in GMEM, 10% FCS (Sigma), NEAA, Na Pyruvate, 0.1 mM ßMercaptoethanol (BMe), Glutamax, and freshly added LIF (1:10,000). 2i-cultured cells were grown on plates coated with Fibronectin, in DMEM:F12/Neurobasal 1:1, KnockOut Serum Replacement, N2, B27, Glutamax, 1µM PD0325901, 3µM CHIR9902, 0.1 mM BMe, and freshly added LIF as above. DuxHA/MERVL-GFP cells were cultured in 2i conditions. siRNAs were purchased from Horizon Discovery (previously Dharmacon) or Sigma (for ‘enzymatically-derived’ esiRNAs). siRNA knockdowns (KDs) were performed for 24hr with the exception of those in Figure 5 which were performed for 72hr. Knockdowns were performed in 6-well plates where 200,000 cells were seeded for 72 hr KDs, and 400,000 for 24 hr KD. 50pmol siRNAs were transfected using RNAiMax Lipofectamine at the time of seeding, and after 48 hrs for 72hr timepoints. Two siRNA controls were used, scrambled (scr) was D-001810-10 and Luciferase (esiLuc) control purchased from Sigma. siSA1 ‘SmartPool’ (SP) was derived from equimolar ratios of commercial siRNAs (D-041989-02, -04, - 05, -06, -07, -08). siSA1 5p was a custom Duplex siRNA sequence (AGGAGCAGGUCGUGGAAGAUU). siSA1 3p was derived from equimolar ratios of commercial siRNAs J-041989-05, -07, -08. esiRNA to SATS was purchased from Sigma as a custom-made product to the entire SATS 5’UTR (mm10 chr9:100,597,794-100,598,109).

### qRT-PCR analysis

Total RNA was isolated using Monarch RNA prep kit (NEB). Reverse transcription was performed on 0.5 µg DNase-treated total RNA using Lunascript RT (NEB) in 20µl reactions. qPCR was performed using 2x SensiFAST SYBR No-ROX kit (Bioline) in 20 µl reactions using 1µl of RT reaction as input and 0.4µM each primer.

### Alkaline Phosphatase (AP) assay and quantification

Cells were seeded in 6 well plates and transfected with siRNAs at the time of plating as above. After 24 hrs, cells were collected for RNA isolation and KD efficiency analyzed by qRT-PCR. Cells from each condition were counted and 1,000 cells per well seeded into a new 6-well plate. Cells were re-transfected after 48 hrs using 5 pmol of siRNAs. Cells were fed every day. Four days after seeding cells at clonal density, the cells were assayed for alkaline phosphatase (AP) expression using StemTAG Alkaline Phosphatase staining kit (Cell Biolabs CBA-300). AP stained cells were imaged in 6-well plates using a M7000 Imaging System (Zeiss) with a 4X objective and a Trans-illumination brightfield light source. For quantification, AP-high and AP-low colonies from each condition were counted. Area occupied by AP-high colonies was also measured using ImageJ, and plotted as fraction of total area of all colonies.

### RACE (Rapid Amplification of cDNA Ends) and PCR mini screen

RACE was performed using GeneRacer kit (RLM RACE, Invitrogen L1500). 2µg of total RNA was used as input. Final products were amplified by nested PCR, using Kapa 2x MasterMix. First PCR was done in a 50µl reaction using 1µl RT as input, 25 cycles. DNA was purified using Qiagen PCR Purification kit, and nested PCR was performed on a tenth of the first PCR for 30 cycles. Viewpoint for 5’RACE was in exon 2 (Fig 3A) or exon 8 (Fig 3B) of Stag1. Viewpoint for 3’RACE was in exon 23 (Fig 3C). RACE primer details can be found in Table S3. PCR products were excised from the gel, A-tailed using Klenow exo- (NEB) and cloned into pCR4-TOPO vector (Invitrogen). At least three clones were sequenced per PCR product. For the PCR Mini-Screen, forward primers at either SATS or canonical 5’ UTR were used with reverse primers either at the end of Stag1 canonical coding sequence, or at the end of coding sequence in intron 25 (see Table S3). PCR was performed using Kapa 2x MasterMix. DNA was excised from the gel, A tailed, and cloned into pCR4-TOPO. At least six clones per PCR product were Sanger-sequenced. Sequences from the PCR Mini-screen were aligned using Minimap2 (2.14-r884) in ‘splice’ mode to ensure long read splice alignment (Fig 3D and S3A).

### PONDR Predictions

Internally disordered regions were predicted using VSL2 predictor at http://www.pondr.com.

### CRISPR-Mediated Stag1 Knock-in Cell Line Generation

The guide RNA targeting Stag1 3’ terminal coding region was designed using Tagin Software (http://tagin.stembio.org) and purchased from IDT. Lyophilised gRNA was rehydrated in RNA duplex buffer (100µM). The single stranded oligodeoxynucleotides (ssODN) encoding mNeonGreen (mNG)-V5-FKBP12^F36V^ and the left and right homology arms was designed using the software tool ChopChop (https://chopchop.cbu.uib.no) and purchased as a High-Copy Amp-resistant plasmid from Twist Bioscience. 2.2µl gRNA (100µM) was mixed with 2.2µl tracrRNA ATTO 550nm (IDT) and annealed together. The RNA duplex was then incubated with 20µg S.p Cas9 Nuclease V3 (IDT) for 10min at room temperature and stored on ice prior to transfection. Linearised KI sequence was mixed with 100% DMSO and denatured at 95°C for 5min. The ssODN was plunged immediately into ice. The RNP complex was mixed with confluent 2i-grown ES cells re-suspended in P3 transfection buffer (Lonza) before being transferred to an electroporation microcuvette well (Lonza). Transfection was performed using a 4D Amaxa electroporator. Post-nucleofection, the cells were seeded into a fibronectin-coated 6 well plate with fresh ESC media. The media was changed daily for four days before being expanded into a T75 flask. Confluent ESC were FACS sorted for GFP+ population (BD FACS Aria Fusion Cell Sorter) and sparsely seeded into 10 cm plates. Clones were manually picked into 96 well plates and expanded for selection by v5 IF, genotyping and Sanger sequencing.

### Dox-inducible Stag1-GFP isoform cell lines

Stag1 isoforms were cloned into pCW57.1 vector (Addgene 41393), modified using Gibson assembly to include an EGFP tag at the 3’end of the Gateway cassette, using Gateway recombination by LR clonase. For primers used to clone the isoforms see Supplementary Table S3. Plasmids were transfected into 2i-grown ESCs using Lipofectamine 3000 and cells grown in Puromycin-supplemented media (1µg/ml) for ten days to make stable lines. Isoform expression was induced using 2µg/ml Doxycycline for 24 hrs, and the population enriched for GFP-positive cells using FACS. For IF experiments, isoforms were induced by adding Dox for 48 hours.

### Protein Lysates, Fractionations and Western blotting

Whole cell lysates (WCL) were collected by lysis in RIPA buffer (150mM NaCl, 1% NP-40 detergent, 0.5% Sodium Deoxycholate, 0.1% SDS, 25mM Tris-HCl pH 7.4, 1mM DTT) and sonicated at 4°C for x5 30 second cycles using Diagenode Bioruptor. Insoluble material was pelleted and the supernatant lysate was quantified using BSA Assay (Thermo Scientific). For cellular fractionations, a cellular ratio of 5×10^6^ cells/80µl buffer was maintained throughout the protocol. Cells were re-suspended in Cell Membrane Lysis Buffer (0.1% Triton X, 10mM HEPES pH 7.9, 10mM KCl, 1.5mM MgCl2, 0.34M sucrose, 10% glycerol, 1mM DTT), incubated on ice for 5min and centrifuged for 5min at 3700rpm to collect the cytoplasmic sample. The pellet was washed and then re-suspended in Nuclear Lysis Buffer (3mM EDTA, 0.2mM EGTA, 1mM DTT) and incubated on ice for 1 hr. Nuclear lysis was aided by sonication with a handheld homogeniser (VWR) for 10sec at 10min intervals. The nucleoplasmic supernatant and chromatin pellet were separated by centrifugation at 9000rpm for 10min at 4°C. The chromatin pellet was re-suspended in 160µl 2X Laemmli Buffer (Bio-Rad). Equal volumes of each fraction were used for Western Blotting (WB). Cytoplasmic and nucleoplasmic protein samples were diluted in 2X Laemmli Buffer and boiled for 5min at 95°C, then loaded on a 4-20% SDS-PAGE gel (Bio-rad) or a 3-8% Tris Acetate gel (Invitrogen). Proteins were wet transferred onto a PDVF membrane (Millipore) and assessed for successful transfer with Ponceau Red (Sigma). The membrane was blocked with 10% milk and incubated with primary antibodies in 1% milk, 0.1% Tween-PBS overnight at 4°C. Membranes were imaged with SuperSignal West Femto Maximum Sensitivity (Thermo) on an ImageQuant.

### Chromatin Co-Immunoprecipitation (co-IP)

Cells were re-suspended in 0.1% NP-40-PBS (1ml/1×10^7^ cells) with 1X Protease Inhibitors (Roche) and 1mM DTT, and centrifuged at 1500rpm for 2min at 4°C. The pellet was re-suspended in Nuclear Lysis Buffer (3mM EDTA, 0.2mM EGTA, 1X Protease Inhibitors, 1mM DTT), vortexed for 30sec before being incubated on a rotator for 30min at 4°C and centrifuged at 6500g for 5min at 4°C to isolate the glassy chromatin pellet. This was re-suspended in High Salt Chromatin Solubilisation Buffer (50mM Tris-HCl pH 7.5, 1.5mM MgCl2, 300mM KCl, 20% glycerol, 1mM EDTA, 0.1% NP-40, 1mM Pefabloc, 1X Protease Inhibitors, 1mM DTT) with Benzonase (Sigma) (6U/1×10^7^) and incubated on rotator for 30min at 4°C. Chromatin was digested with 3x 10sec sonication at 30% intensity with a Vibra-Cell probe. The supernatant was collected by centrifugation at 1300rpm for 30min at 4°C, and then diluted to 200mM KCl concentration with no KCL buffer. 30µl of Dynabeads (Invitrogen) were used per co-IP. Beads were washed 2x in 200mM KCl IP Buffer, re-suspended in IP Buffer with 10µg of the IP antibody, or an IgG-containing serum to match the species of the IP antibody and placed on rotator for 5h at 4°C. Beads were washed 3x in IP buffer and then incubated in 1mg chromatin lysate on a rotator overnight at 4°C. The beads were washed, re-suspended in 2X Laemmli Buffer (Bio-Rad), boiled for 10min at 95°C and used for WB as above.

### Immunofluorescence and Microscopy

ESCs were cultured on fibronectin or gelatin-coated cover glass in 6-well plates. Cells were fixed in 4% Paraformaldehyde for 5min and incubated in 0.1% Triton X-PBS for 10min before being washed and blocked in 10% FCS-PBS for 20min. Primary antibodies were diluted in 10% FCS, 0.1% Saponin (Sigma) and incubated overnight at 4°C. The next day, the cells were incubated with an Alexa fluorophore-conjugated secondary antibody diluted in 10% FCS, 0.1% Saponin for 1 hr at room temperature, washed and mounted on cover slides with ProLong Diamond Antifade Mountant with DAPI (Invitrogen). Z-stacks imaging of fixed cells was done using a LSM 880 confocal microscope (Zeiss) with a 63X oil objective. Analysis was performed using Imaris 9.6 (Oxford instruments). Live cell imaging was performed using a 3i Spinning Disc confocal microscope (Zeiss). Stag1-mNG-V5-FKBP12^F36V^ cells were seeded in an 8-chambered coverglass (Lab-Tek II) and DMSO or dTAG (500nM) were added for 24hr before imaging. Directly prior to imaging, cells were incubated with Hoechst 33342 (BD Pharmingen) for 45min, and then replaced with fresh 2i ESC media. Cells were imaged as confocal Z-stacks using DAPI and GFP lasers with a 63X objective and 1.4 Numerical Aperture.

### Antibodies used in this study

| Protein | Catalogue No. | Company | Figure references |
| --- | --- | --- | --- |
| Stag1/SA1, N-term epitope | ab4455 | Abcam | 1B, C, I, S1C, K, 2C, S2C, E, 3J, S3G, 4C, F, 5J |
| Stag1/SA1, C-term epitope | ab4457 | Abcam | 2F, S5A, 3I |
| Stag2/SA2 | A300-158A | Bethyl | 1B, S1C |
| Smc3 | ab9263 | Abcam | 1B, 2C |
| Nanog | ab70482 | Abcam | 1E, S1F |
| Tubulin (Tub) | T5168 | Sigma | 1C, 1I, S2E, 4C, S6A |
| Actin | Mab8929 | Novus | S1C |
| H3 | ab1791 | Abcam | 1C |
| v5 | 14-6796-82 | Invitrogen | 3K |
| HP1a | 2616 | Cell Signalling | 2C, S2B, C |
| Nucleolin (Ncl) | ab22758 | Abcam | 2C, 5J, 6A, S6A |
| POLR2 | MMS-128P | Covance | 3K, L |
| H3K9me3 | ab8898 | Abcam | 2F, I, S2E, 5A, S5A |
| H3K4me3 | ab8580 | Abcam | S2E |
| Alexa488-anti-GFP (GFP) | A-21311 | ThermoFisher | 2I, S2A, B, 5A |
| Trim28 | MA1-2023 | ThermoFisher | 5J |

### Nascent transcription and translation analysis

For nascent transcription analysis, we used the Click-iT® RNA Alexa Fluor® 488 HCS Assay (Invitrogen C10327). ES cells were labelled with 1mM EU for 45min at 37C in fresh ES media. Cells were fixed in solution or onto coverslips with 3.7% paraformaldehyde and permeabilised with 0.5% Triton-X solution. Cells were incubated with the Click-iT reaction cocktail for 30min. Cells were then either processed further for Immunofluorescence as per methods described above (directly to the blocking step) or analysed by flow cytometry on a BD Fortessa X20. For the Nascent translation analysis, Click-iT™ HPG Alexa Fluor™ 594 Protein Synthesis Assay Kit (Invitrogen C10429) was used. Cells were pre-incubated in Methionine-free media for 30 min in the 37C incubator before addition of L-homopropargylglycine (HPG) at 50µM. Cells were incubated with HPG for 30 min, then collected, fixed, permeabilized, and stained using Click-It reaction in low retention tubes. HPG incorporation was measured by Flow Cytometry. FACS analysis (in Figures 5,6) was done with FloJo software (version 10.7.1).

### Next generation Sequencing and Analysis

Genomic data generated in this study (RNA-seq, PacBio-seq and UMI4C-seq) was submitted to GEO with the Accession GSE160390.

### RNA sequencing (RNA-seq) library preparation and sequencing

ESCs were treated for 24hrs with siRNA pools to Stag1 (SA1) and two sets of control siRNAs, scrambled (SCR) and Luciferase (Luc). There are three replicate sets for SP KD and two for the siRNA pools (SATS, 3p, 5p). Total RNA was isolated using NEB Monarch RNA prep kit. 1µg of total RNA was rRNA-depleted using NEBNext rRNA depletion kit (Human/Mouse/Rat). Libraries were prepared from 10-50ng rRNA-depleted total RNA, depending on availability of material, using NEBNext Ultra II directional RNAseq kit according to manufacturer’s instructions using 8 cycles of PCR. All ESC FCS libraries were rRNA depleted and only the ESC 2i libraries were PolyA-enriched before library prep. Two rounds of PolyA+ enrichment were performed. RNA-seq libraries were sequenced on the Illumina HiSeq3000 platform, 75bp paired-end or single-end reads. Reads were quality controlled using FASTQC. RNA-seq data was processed using the RNA-seq Nextflow pipeline (v19.01.0), with the following parameters –aligner hisat2 – genome mm10, with –reverse_stranded specified for paired-end samples. FeatureCounts output was parsed through edgeR (v3.16.5) and DESeq2 (v1.14.1) to generate normalised expression counts. The normalised counts for RNAseq (Figure 1) were calculated in edgeR. Low expressed genes were removed (rowSum cpm <2 across SCR and SA1SP replicates), normalisation factors were calculated using calcNormFactors and dispersions estimated using estimateDisp. The edgeR volcano plot statistics were calculated using the exactTest and topTags functions. To generate the normalised counts for RNAseq experiments required to calculate the log2FC GSEA ranked lists, the FeatureCounts output for all experiments was combined into a single table and read into DESeq2. A DESeq2 object was built using the function DESeqDataSetFromMatrix and estimation of size factors and dispersions were calculated using the DEseq function. Normalised counts were calculated using the ‘counts’ function. Low expressed genes (rowSum normalised count <10 across all samples) were removed.

### GSEA

Broad Institute GSEAPreranked (v4.0.3) was used to determine the enrichment of curated genesets within our RNA-seq data. For each sample a ranked list was generated with genes ranked in descending order by their log2FC value using normalised expression scores from DEseq2. Log2FC per gene was calculated between the KD and its respective SCR using the following calculation: Log2(normalised_counts KD +1) - log2(normalised_counts SCR +1). In the case of experiments with multiple KD replicates, the average log2 normalised count was used. Three gene sets were assayed in this study, ‘naïve pluripotency’, ‘primed pluripotency’ and ‘2C signatures’. The naïve and primed pluripotency gene sets were curated in-house from Fidalgo M et al. (CSC, 2016) where genes were selected if they had >2 fold change. The naïve and primed gene sets contained 661 and 580 genes respectively. The 2C signatures gene set (147 genes) was obtained from Percharde M et al. (Cell, 2018). Gene sets were classed as having significant enrichment if the p-value was <0.05 and the normalised enrichment score (NES) exceeded +/- 1.

### VAST-TOOLS

VAST-TOOLS was used to generate Percent Spliced In (PSI) scores, a statistic which represents how often a particular exon is spliced into a transcript using the ratio between reads which include and exclude said exon. Paired-end RNA-seq datasets were submitted to VAST-TOOLS (v2.1.3) using the Mmu genome (Tapial J et al, Gen Res 2017). Briefly, reads are split into 50nt words with a 25nt sliding window. The 50nt words are aligned to a reference genome using Bowtie to obtain unmapped reads. These unmapped reads are then aligned to a set of predefined exon-exon junction (EJJ) libraries allowing for the quantification of alternative exon events. The output was further interrogated using a script which searches all hypothetical EEJ combinations between potential donors and acceptors within Stag1. PSI scores could be obtained providing there was at least a single read within our RNAseq data that supported one of these potential events. Some datasets were combined to have enough reads for the analysis. See Table S1 for PSI values and names of RNA-seq libraries used for analysis in Fig. 3e, S4b.

### Quantifying sectioned Stag1

Stag1 was split into 5 sections; SATS, e1-e8, e12-e19, e20-e25, e26-e34. Using Kallisto (v0.46.1), raw RNAseq reads were used to quantify each section of Stag1. Kallisto was run in quant mode, using the –rf-stranded parameter, outputting a TPM per Stag1 section. A line plot was generated showing TPM in relative to UT.

### PacBio library, sequencing and analysis

ES cells were cultured in naïve 2i conditions and PolyA-enriched mRNAs were hybridized to a custom Biotinylated oligonucleoltide probe set. Post-capture, mRNAs were amplified using the Clontech SMARTer PCR cDNA Synthesis Kit with 9 cycles and used in the SMRTbell library prep according to manufacturers instructions. The library was sequenced on the SMRTseq 2000 platform. PacBio reads were processed through the SMRTLINK v8.0.0 IsoSeq3 pipeline. 403,995 Circular consensus sequences (CCS) were generated using default parameters (--minPasses = 1, --min-rq = 0.8, CCS Polish = No). Further refining through lima (removal of adapters and correct orientation of sequences), poly-A trimming and concatemer removal resulted in 265,106 full length non-chimeric (FLNC) reads. FLNC reads were aligned to the mm10 genome using Minimap2 with the following parameters (-ax splice, -uf, -k14).

### ChIP-seq Analysis

Previously published Stag1 Chromatin Immunoprecipitation-sequencing (ChIP-seq) datasets from ES 2i cells (GSE126659, only Replicate 1 and 2 libraries) were trimmed using trim_galore and aligned to mm10 using bowtie2. Peak detection was performed with MACS2 using uniquely reads (MAPQ≥2). Peaks were overlapped with genomic features in a hierarchical manner (promoters > exons > repeats > introns > intergenic), and overlap frequency was compared with a randomly shuffled version of the peaks. To identify repeat families enriched for STAG1 peaks, a previously described pipeline was used (Deniz O et al. Nat Comm, 2020) that compares family-levels overlap frequency with that observed in 1,000 permutations of random peak shuffling. Coverage profiles across specific TE families were generated using HOMER and including multi-mapping reads (MAPQ<2).

### UMI-4C library preparation

1×10^7^ cells were fixed at RT for 10min in 1% formaldehyde and fixation was quenched with 0.125M Glycine for 5min. Cells were then lysed on ice in 10ml Lysis Buffer (10mM NaCl, 10mM Tris-HCl pH 8.0, 0.25% NP40, protease inhibitor) for 30min, followed by 10 strokes of douncing using a tight pestle. Nuclei were pelleted, 8min 700 rcf, washed in 1ml 1.2X DpnII buffer in Protein LoBind tubes (Eppendorf) and resuspended in 500 µl 1.2X DpnII buffer. 15ul of 10% SDS was added and incubated for 1hr at 37°C shaking at 650 rcf. 50ul of 20% TritonX was added to quench the SDS and incubated for 15 min at 37°C with shaking. 750U of DpnII was added and incubated overnight at 37**°C** with interval shaking. The next morning, nuclei were pelleted at 4°C by 650 rcf for 5 min and resuspended in 500µl 1X DpnII buffer. 500U DpnII was added and incubated for an additional four hours. The nuclei were washed twice in 100 µl of 1X T4 Ligase Buffer and resuspended in 200 µl Ligase Buffer. 6ul of T4 DNA Ligase was added and incubated for 3hr at 16°C. Nuclei were then pelleted, resuspended in 200 µl 1x fresh Ligase Buffer, 6µl of T4 DNA Ligase added, and incubated overnight at 16°C. Samples were treated with 20µl of ProtK (NEB Molecular Biology Grade), incubated for 3 hrs at 55°C and 5 hrs at 65°C to reverse crosslinks. Samples were treated with RNase A (PureLink, Invitrogen) for 1 hr at 37°C and DNA was extracted and precipitated overnight. For library preparation, 3×5µg of ligated DNA was sonicated using Covaris (10% duty cycle, intensity 5, cycle burst 200, 70sec). Samples were end-repaired using DNA PolII Klenow Large Fragment (NEB), A-tailed using Klenow (exo-) (NEB), and Illumina indexed adapters ligated using Quick DNA Ligase (NEB). Reactions were denatured at 95°C for 3 min, placed on ice, and purified using 1.2X SizeSelect AmpPure beads to recover ssDNA. Libraries were amplified using GoTaq (Promega), with 20 cycles for PCR1 and 15 cycles for nested PCR2 on 50% material from 1^st^ PCR. For custom UMI bait sequences, see Table S3.

### Hi-C and UMI-4C-seq analysis

Hi-C libraries were analysed as previously described (Barrington 2019). UMI-4C tracks were processed using the ‘umi4cPackage’ pipeline (v0.0.0.9000) (Schwartzman, O et al. Nat Meth 2017). Briefly, raw reads are parsed through the UMI-4C pipeline, those reads containing the bait and padding sequence are retained and de-multiplexed. Reads lacking the padding sequence are considered non-specific and are removed from further analysis. Retained reads are split based on a match to the restriction enzyme sequence to create a segmented fastq file. The first 10 bases of read 2 are extracted and attached to the segments derived from each read pair. Mapping to mm10 is done with Bowtie2. Read pairs that have reverse complement segments are mapped to a restriction fragment ID, with the fragment ID, strand and distance from each end represented within a fragment-chain table. UMI filtering is used to determine the number of molecules supporting each ligation event. The resulting UMI-4C tracks are then imported into R, and data from multiple bait replicates can be merged by summing the molecule counts per ligated fragment, at which point contact intensity profiles and domainograms around the viewpoint can be generated (see Figure 3). The contact intensity profile represents the mean number of ligations within a genomic window, with the resolution of the contact intensity profile being determined by the window size (set to 15 here). The domainogram reports the mean contact per fend at a series of window sizes, a stacked representation of contact intensity values in increasing window sizes from 10 to 300 fragment ends, their colour can be used to identify peak locations. ES and NSC contact profiles were compared after normalisation to correct for bias (see Schwartzman et al for further details). For the compared profiles, the total molecule count for restriction fragment ends for each are calculated at three ranges around the viewpoint. One profile is selected as a reference and the second is scaled to the first using the ratio in total molecule counts between the two profiles as the scaling factor. Below the contact profile is the profile resolution indicator, which shows the number of fends required to include at least 15 UMI molecules. The darker the colour, the larger the window size required. The domainogram at the bottom represents the log2 ratio between the domainogram values of the compared profiles and highlights locations where ESC has more contacts than NSC or vice versa.

## REFERENCES

1. Horsfield, J. A. et al. Cohesin-dependent regulation of Runx genes. Development 134, 2639–2649 (2007).

2. Viny, A. D. et al. Cohesin Members Stag1 and Stag2 Display Distinct Roles in Chromatin Accessibility and Topological Control of HSC Self-Renewal and Differentiation. Cell Stem Cell 25, 682–696.e8 (2019).

3. Leiserson, M. D. M. et al. Pan-cancer network analysis identifies combinations of rare somatic mutations across pathways and protein complexes. Nat. Genet. 47, 106–114 (2015).

4. Romero-Pérez, L., Surdez, D., Brunet, E., Delattre, O. & Grünewald, T. G. P. STAG Mutations in Cancer. Trends Cancer 5, 506–520 (2019).

5. Cuartero, S. et al. Control of inducible gene expression links cohesin to hematopoietic progenitor self-renewal and differentiation. Nat Immunol 19, 932–941 (2018).

6. Kline, A. D. et al. Diagnosis and management of Cornelia de Lange syndrome: first international consensus statement. Nat. Rev. Genet. 19, 649–666 (2018).

7. Hadjur, S. et al. Cohesins form chromosomal cis-interactions at the developmentally regulated IFNG locus. Nature 460, 410–413 (2009).

8. Phillips-Cremins, J. E. et al. Architectural protein subclasses shape 3D organization of genomes during lineage commitment. Cell 153, 1281–1295 (2013).

9. Wendt, K. S. et al. Cohesin mediates transcriptional insulation by CCCTC-binding factor. Nature 451, 796–801 (2008).

10. Parelho, V. et al. Cohesins functionally associate with CTCF on mammalian chromosome arms. Cell 132, 422–433 (2008).

11. Mishiro, T. & Tsutsumi, S. Architectural roles of multiple chromatin insulators at the human apolipoprotein gene cluster. EMBO J. 28, 1234–1245 (2009).

12. Kagey, M. H. et al. Mediator and cohesin connect gene expression and chromatin architecture. Nature 467, 430–435 (2010).

13. Misulovin, Z. et al. Association of cohesin and Nipped-B with transcriptionally active regions of the Drosophila melanogaster genome. Chromosoma 117, 89–102 (2007).

14. Vietri Rudan, M. et al. Comparative Hi-C reveals that CTCF underlies evolution of chromosomal domain architecture. CellReports 10, 1297–1309 (2015).

15. Rao, S. S. P. et al. A 3D map of the human genome at kilobase resolution reveals principles of chromatin looping. Cell 159, 1665–1680 (2014).

16. Sofueva, S. et al. Cohesin-mediated interactions organize chromosomal domain architecture. EMBO J. 32, 3119–3129 (2013).

17. Zuin, J. et al. Cohesin and CTCF differentially affect chromatin architecture and gene expression in human cells. Proc. Natl. Acad. Sci. U.S.A. 111, 996–1001 (2014).

18. Rao, S. S. P. et al. Cohesin Loss Eliminates All Loop Domains. Cell 171, 305– 309.e24 (2017).

19. Seitan, V. C. et al. Cohesin-based chromatin interactions enable regulated gene expression within preexisting architectural compartments. Genome Res. 23, 2066–2077 (2013).

20. Schwarzer, W. et al. Two independent modes of chromatin organization revealed by cohesin removal. Nature 551, 51–56 (2017).

21. Wutz, G. et al. Topologically associating domains and chromatin loops depend on cohesin and are regulated by CTCF, WAPL, and PDS5 proteins. EMBO J. 36, 3573–3599 (2017).

22. Haarhuis, J. H. I. et al. The Cohesin Release Factor WAPL Restricts Chromatin Loop Extension. Cell 169, 693–707.e14 (2017).

23. Lehalle, D. et al. STAG1 mutations cause a novel cohesinopathy characterised by unspecific syndromic intellectual disability. J Med Genet 54, 479–488 (2017).

24. Soardi, F. C. et al. Familial STAG2 germline mutation defines a new human cohesinopathy. NPJ Genom Med 2, 7–11 (2017).

25. Yuan, B. et al. Clinical exome sequencing reveals locus heterogeneity and phenotypic variability of cohesinopathies. Genet Med 21, 663–675 (2019).

26. Cuadrado, A. et al. Specific Contributions of Cohesin-SA1 and Cohesin-SA2 to TADs and Polycomb Domains in Embryonic Stem Cells. Cell Rep 27, 3500–3510.e4 (2019).

27. Hara, K. et al. Structure of cohesin subcomplex pinpoints direct shugoshin-Wapl antagonism in centromeric cohesion. Nature Publishing Group 21, 864–870 (2014).

28. Xiao, T., Wallace, J. & Felsenfeld, G. Specific sites in the C terminus of CTCF interact with the SA2 subunit of the cohesin complex and are required for cohesin-dependent insulation activity. Mol. Cell. Biol. 31, 2174–2183 (2011).

29. Li, Y. et al. The structural basis for cohesin-CTCF-anchored loops. Nature 578, 472–476 (2020).

30. Orgil, O. et al. A conserved domain in the scc3 subunit of cohesin mediates the interaction with both mcd1 and the cohesin loader complex. PLoS Genet. 11, e1005036 (2015).

31. Canudas, S. & Smith, S. Differential regulation of telomere and centromere cohesion by the Scc3 homologues SA1 and SA2, respectively, in human cells. J Cell Biol 187, 165–173 (2009).

32. Kojic, A. et al. Distinct roles of cohesin-SA1 and cohesin-SA2 in 3D chromosome organization. Nature Publishing Group 25, 496–504 (2018).

33. Remeseiro, S. et al. Cohesin-SA1 deficiency drives aneuploidy and tumourigenesis in mice due to impaired replication of telomeres. EMBO J. 31, 2076–2089 (2012).

34. Winters, T., McNicoll, F. & Jessberger, R. Meiotic cohesin STAG3 is required for chromosome axis formation and sister chromatid cohesion. EMBO J. 33, 1256–1270 (2014).

35. Bisht, K. K., Daniloski, Z. & Smith, S. SA1 binds directly to DNA through its unique AT-hook to promote sister chromatid cohesion at telomeres. J. Cell. Sci. 126, 3493–3503 (2013).

36. Boisvert, F.-M., van Koningsbruggen, S., Navascués, J. & Lamond, A. I. The multifunctional nucleolus. Nat. Rev. Mol. Cell Biol. 8, 574–585 (2007).

37. Feric, M. et al. Coexisting Liquid Phases Underlie Nucleolar Subcompartments. Cell 165, 1686–1697 (2016).

38. Yao, R.-W. et al. Nascent Pre-rRNA Sorting via Phase Separation Drives the Assembly of Dense Fibrillar Components in the Human Nucleolus. Molecular Cell 76, 767–783.e11 (2019).

39. Padeken, J. & Heun, P. Nucleolus and nuclear periphery: velcro for heterochromatin. Curr. Opin. Cell Biol. 28, 54–60 (2014).

40. Kresoja-Rakic, J. & Santoro, R. Nucleolus and rRNA Gene Chromatin in Early Embryo Development. Trends Genet. 35, 868–879 (2019).

41. Aguirre-Lavin, T. et al. 3D-FISH analysis of embryonic nuclei in mouse highlights several abrupt changes of nuclear organization during preimplantation development. BMC Dev Biol 12, 30–20 (2012).

42. Fulka, H., Rychtarova, J. & Loi, P. The nucleolus-like and precursor bodies of mammalian oocytes and embryos and their possible role in post-fertilization centromere remodelling. Biochemical Society Transactions 48, 581–593 (2020).

43. Ishiuchi, T. et al. Early embryonic-like cells are induced by downregulating replication-dependent chromatin assembly. Nature Publishing Group 22, 662–671 (2015).

44. Xie, S. Q. et al. Nucleolar-based Dux repression is essential for embryonic two-cell stage exit. Genes Dev. 36, 331–347 (2022).

45. Németh, A. et al. Initial genomics of the human nucleolus. PLoS Genet. 6, e1000889 (2010).

46. Gupta, S. & Santoro, R. Regulation and Roles of the Nucleolus in Embryonic Stem Cells: From Ribosome Biogenesis to Genome Organization. Stem Cell Reports 15, 1206–1219 (2020).

47. Grow, E. J. et al. p53 convergently activates Dux/DUX4 in embryonic stem cells and in facioscapulohumeral muscular dystrophy cell models. Nat. Genet. 53, 1207–1220 (2021).

48. Sun, Z. et al. LIN28 coordinately promotes nucleolar/ribosomal functions and represses the 2C-like transcriptional program in pluripotent stem cells. Protein Cell 1–23 (2021). doi:10.1007/s13238-021-00864-5

49. Laloraya, S., Guacci, V. & Koshland, D. Chromosomal addresses of the cohesin component Mcd1p. Journal of Cell Biology 151, 1047–1056 (2000).

50. Harris, B. et al. Cohesion promotes nucleolar structure and function. Mol Biol Cell 25, 337–346 (2014).

51. Mootha, V. K. et al. PGC-1alpha-responsive genes involved in oxidative phosphorylation are coordinately downregulated in human diabetes. Nat. Genet. 34, 267–273 (2003).

52. Subramanian, A. et al. Gene set enrichment analysis: a knowledge-based approach for interpreting genome-wide expression profiles. Proc Natl Acad Sci USA 102, 15545–15550 (2005).

53. Nabet, B. et al. The dTAG system for immediate and target-specific protein degradation. Nature Chemical Biology 14, 1–16 (2018).

54. Quinodoz, S. A. et al. Higher-Order Inter-chromosomal Hubs Shape 3D Genome Organization in the Nucleus. Cell 174, 744–757.e24 (2018).

55. Deniz, Ö. et al. Endogenous retroviruses are a source of enhancers with oncogenic potential in acute myeloid leukaemia. Nature Communications 11, 3506–14 (2020).

56. Percharde, M. et al. A LINE1-Nucleolin Partnership Regulates Early Development and ESC Identity. Cell 174, 391–405.e19 (2018).

57. Hackett, J. A., Kobayashi, T., Dietmann, S. & Surani, M. A. Activation of Lineage Regulators and Transposable Elements across a Pluripotent Spectrum. Stem Cell Reports 8, 1645–1658 (2017).

58. Dixon, J. R. et al. Topological domains in mammalian genomes identified by analysis of chromatin interactions. Nature 485, 376–380 (2012).

59. Schwalie, P. C. et al. Co-binding by YY1 identifies the transcriptionally active, highly conserved set of CTCF-bound regions in primate genomes. Genome Biol. 14, R148–15 (2013).

60. Rowe, H. M. et al. KAP1 controls endogenous retroviruses in embryonic stem cells. Nature 463, 237–240 (2010).

61. Meshorer, E. et al. Hyperdynamic plasticity of chromatin proteins in pluripotent embryonic stem cells. Dev. Cell 10, 105–116 (2006).

62. Feng, G. et al. Ubiquitously expressed genes participate in cell-specific functions via alternative promoter usage. EMBO Rep. 17, 1304–1313 (2016).

63. Tapial, J. et al. An atlas of alternative splicing profiles and functional associations reveals new regulatory programs and genes that simultaneously express multiple major isoforms. Genome Res. 27, 1759–1768 (2017).

64. Barrington, C., Georgopoulou, D., Nature, D. P.2019. Enhancer accessibility and CTCF occupancy underlie asymmetric TAD architecture and cell type specific genome topology. nature.com doi:10.1038/s41467-019-10725-9

65. Hendrickson, P. G. et al. Conserved roles of mouse DUX and human DUX4 in activating cleavage-stage genes and MERVL/HERVL retrotransposons. Nat. Genet. 49, 925–934 (2017).

66. Macfarlan, T. S. et al. Embryonic stem cell potency fluctuates with endogenous retrovirus activity. Nature 487, 57–63 (2012).

67. Eckersley-Maslin, M. A. et al. MERVL/Zscan4 Network Activation Results in Transient Genome-wide DNA Demethylation of mESCs. Cell Rep 17, 179–192 (2016).

68. Remeseiro, S., Cuadrado, A., López, G. G., Pisano, D. G. & Losada, A. A unique role of cohesin-SA1 in gene regulation and development. EMBO J. 31, 2090–2102 (2012).

69. Borsos, M. & Torres-Padilla, M.-E. Building up the nucleus: nuclear organization in the establishment of totipotency and pluripotency during mammalian development. Genes Dev. 30, 611–621 (2016).

