## Supplemental File for "The N-terminus of Stag1 is required to repress the 2C program by maintaining rRNA expression and nucleolar integrity"

**Supplementary Figure 1. Stag1 is required for pluripotency in mESCs.**

a) Cartoon of the cohesin complex including the core trimer subunits of Smc1a, Smc3 and Rad21 complexed with either Stag1 or Stag2.

b) Relative expression of Stag1 and Stag2 mRNA by qRT-PCR in 2i- (naïve) or FCS-grown mESC, EpiLCs and MEFs. Data is represented as mean ± SEM of two independent experiments and relative to Actin control expression.

c) WCL from naïve (2i) mESC and EpiLCs, sorted for cells in the G1 phase and analysed by WB for levels of SA1 and SA2. Actin serves as a loading control.

d) Relative expression of Stag1 mRNA by qRT-PCR in FCS- (left panel, n=20) or 2i-grown (right panel, n=19) mESCs upon treatment with si scr or si SA1, smartpool (SP). Whiskers and boxes indicate all and 50% of values respectively. Central line represents the median.

e) WB analysis of SA1 levels in WCL, cytoplasmic and chromatin fractions upon treatment with scrambled siRNAs (si Scr) or SA1 siRNAs (siSA1) for 24hr in naïve mESCs. Data as in Fig. 1c, but also including the cytoplasmic fraction. Tubulin (Tub) and H3 serve as fractionation and loading controls.

f) Cell cycle analysis of Hoechst-stained 2i mESC after treatment with si scr or siSA1 siRNAs for 24hrs. Shown are the percentages of cells in G1 or G2 phases. These are the same cells that were used for the RNA-sequencing experiments shown in Figure 1.

g) Left, Relative expression of Nanog mRNA by qRT-PCR in FCS-grown mESCs upon treatment with si scr or siSA1. Quantification and statistics as before.

h) Enrichment score (ES) plots from GSEA using the naïve or primed gene sets as in Figure 1 and RNA-seq data from two siSA1 treated mESC replicates. The third replicate is shown in the main Figure.

i, j) Cartoon of the CRISPR/Cas9 targeting strategy to introduce a NeonGreen-v5-FKBP tag to the C-terminus of endogenous Stag1. FOR and REV indicate primers used for genotyping, shown in j). The leftmost NG/NG homozygous sample represents the 'B1 clone' used throughout the manuscript.

k) Representative confocal images of neon-green in $SA1^{NG\text{-}FKBP}$ mESC (clone B1) treated with DMSO or dTAG.

**Supplementary Figure 2. Stag1 is localised to and impacts both euchromatin and heterochromatin compartments.**

a) Left, representative confocal images of IF to SA1 and H3K9me3 in siRNA-treated mESC counterstained with DAPI. Right, Imaris quantification of the volume of H3K9me3 foci from siRNA treated mESC. Quantifications and statistical analysis were done as above. Data is from two independent biological replicates with n>50/condition.

b) Left, representative confocal images of IF to GFP and H3K9me3 in mESCs expressing a dox-inducible GFP-tagged full-length SA1 (SA1-FL) and counterstained with DAPI. Right, Imaris quantification of the volume of H3K9me3 foci from dox-inducible mESCs. Quantifications and statistical analysis were done as above. Data is from two independent biological replicates with n>40/condition.

c) Percentage of SA1 ChIP-seq peaks in mESCs (unique reads) at promoters, exons, transposable element repeats, introns and intergenic sequences.

d) Analysis of CTCF motifs contained within selected repeat elements and the percentage of Stag1 binding. NB. The majority of elements contain both a CTCF motif and Stag1.

e) SA1 ChIP-seq (unique and multimapping reads) aligned to additional full-length repeat elements. Two SA1 ChIP replicates are shown in blue alongside the INPUT in grey.

f) Relative expression of Stag1, LINE1-T and pre-rRNA by qRT-PCR in siRNA-treated mESCs. Shown are total RNA levels. Data is represented as mean ± SEM and statistical analysis as before. Data is from three independent experiments.

Representative confocal images of MFI of Nucleolin assessed by IF in g) $SA1^{NG\text{-}FKBP}$ mESC (clone B1) treated with DMSO or dTAG (two different cells shown) and h) siRNA-treated mESC and counterstained with DAPI. Quantification of these is shown in Fig. 2k.

**Supplementary Figure 3. Transcription-Regulatory Control of Stag1 in mESCs.**

a) Aligned Stag1 transcript variants identified from 5'RACE in Fig. 3a. Arrows refer to the bands on the RACE gels which were cloned and sequenced. *NB,* the diversity of skipping events that all result in a functional loss of the 5' end of Stag1.

b) Over-exposure of the 5'RACE gel shown in Fig. 3a (right) to better show the small RACE products (blue arrows).

c) Close-up of the 3' RACE sequence that identified a new alternative TTS in intron 25 (sequence shown in dark blue).

d) Initial PCR screen in naïve mESCs and MEFs using various combinations of forward (5') primers (SATS, canonical TSS, Alt exon 1 TSS) and reverse (3') primers (canonical TTS, Alt intron 25 TTS). *NB.* SATS is only expressed in ESC; canonical, full-length Stag1 is more expressed in ESC compared to MEFs; and the alternative intron 25 TTS is most often expressed with a canonical TSS.

e) Full-length transcripts sequenced on the PacBio platform. Including many isoforms already discovered using RACE and PCR cloning methods above.

f) Percent Spliced In (PSI) calculations based on VAST-Tools analysis of RNA-seq from multiple 2i (blue) and FCS (red) datasets (see Methods for details of libraries). Data are shown relative to Neural Stem cell (NSC) frequencies, highlighting the events that are ESC-specific.

g) Top, cartoon depicting functional domains within Stag1 protein, including the AT-hook (aa 3-58); Stromalin conserved domain (SCD, aa 296-381) and the C-terminus. Middle, the predicted Stag1 protein isoforms based on transcript analysis with estimated sizes for each isoform and colour coded according to the analysis in d). Purple boxes in the 105kDa and 90kDa isoforms represent retained introns. Bottom, PONDR (Predictor of Natural Disordered Regions) analysis of SA1 protein using VSL2 predictor at http://www.pondr.com showing consecutive stretches of disordered regions corresponding to the N- and C-terminus of SA1 in its full-length (FL), N-terminal delta (ΔN) and C-terminal delta (ΔC) isoform groups.

h) Chromatin Immunoprecipitation of endogenous SA1 in mESCs and EpiLCs. IgG was used as a control. *NB.* Both canonical and SA1 isoforms reduce in levels upon differentiation.

**Supplementary Figure 4. Genome topology at the Stag1 locus.**

Hi-C contact maps in naïve mESC and NSC of the 900kb region on chromosome 9 containing the Stag1 topologically associated domain (TAD). TADs are denoted with a vertical line and as repressed (orange) or active (blue). Shown also are tracks for Genes, Nanog and CTCF ChIP-seq as well as a track indicating the directionality of CTCF binding sites (red, forward; blue, reverse). Aligned to the Gene track are also the SA1 transcripts discovered above where red represents the untranslated regions and blue the coding body. UMI-4C-seq viewpoints are positioned to the leftmost CTCF site ('CTCF bait', vertical green arrow on ChIP track) and to a Nanog site 40 kb upstream of the Stag1 canonical TSS ('Nanog bait', vertical purple arrow on ChIP track). For each bait, UMI information for each cell type is shown as well as the comparative plots where red represents an enrichment of contacts in ESC compared to NSC.

**Supplementary Figure 5. Fluctuations in Stag1 isoforms skews cell fates.**

a) Relative expression of Stag1 mRNA by qRT-PCR in FCS- (n=7) or 2i-grown (n=6) mESC upon si scr or the si SA1 panel. Quantifications as before. NB, all siRNAs knockdown Stag1 to a similar extent with the exception of esiRNA SATS which reduced Stag1 by 40%.

b) Left, relative expression of Nanog mRNA by qRT-PCR in FCS-grown mESCs upon si scr or the si SA1 panel (n=13). Quantifications as before. *NB,* the modest, but different influence of the 5p and the 3p KDs on Nanog levels. Right, WB analysis of Nanog levels in siRNA treated mESC WCL. Tubulin serves as a loading control. The percentage of knockdown (KD) of Nanog protein signal normalised to Tubulin is shown.

c) Enrichment score (ES) plots from GSEA using the naïve or primed gene sets as in Fig. 1e and RNA-seq data from mESCs treated with the siRNAs to SATS TSS, 3p and 5p.

d) AP+ colonies in mESCs (purple), as a percentage of all colonies (pink and purple) treated with the siRNA panel. Data are the average of two independent biological replicates. See also Fig. 4f.

e) Global analysis of nascent transcription by measuring EU-488 incorporation using Flow cytometry. Left, representative Flo-Jo analysis of EU incorporation in mESCs treated with the siRNA KD panel and controls. Right, quantification of the change in EU incorporation relative to si scr treated cells. Data are represented as the mean +/- SEM and are from two biological replicates. Statistical analysis using two-tailed t-test.

f) Global analysis of nascent translation by measuring HPG incorporation using Flow cytometry. Shown are representative Flo-Jo analysis of HPG incorporation in mESC treated with the siRNA KD panel. Quantifications of the data can be found in Fig. 5e.

g) Relative expression of Rpl3 and Rps9 mRNA by qRT-PCR in mESC upon si scr or the si SA1 panel. Data are represented as the mean +/- SEM and are from three biological replicates. Statistical analysis using two-tailed t-test.

**Supplementary Figure 6. Loss of the Stag1 N-terminus leads to conversion to totipotency.**

a) Relative expression of 2C related genes by qRT-PCR in 2i-grown mESC after treatment with si scr or the si SA1 panel. Data are represented as mean +/- SEM from n=5 biological replicates.

b) Enrichment score (ES) plots from GSEA using 2C gene sets as in Fig. 6e and the biological replicate RNA-seq data from the different siRNA treated mESC samples.

**Supplementary Figure 1.** *Pezic et al.*

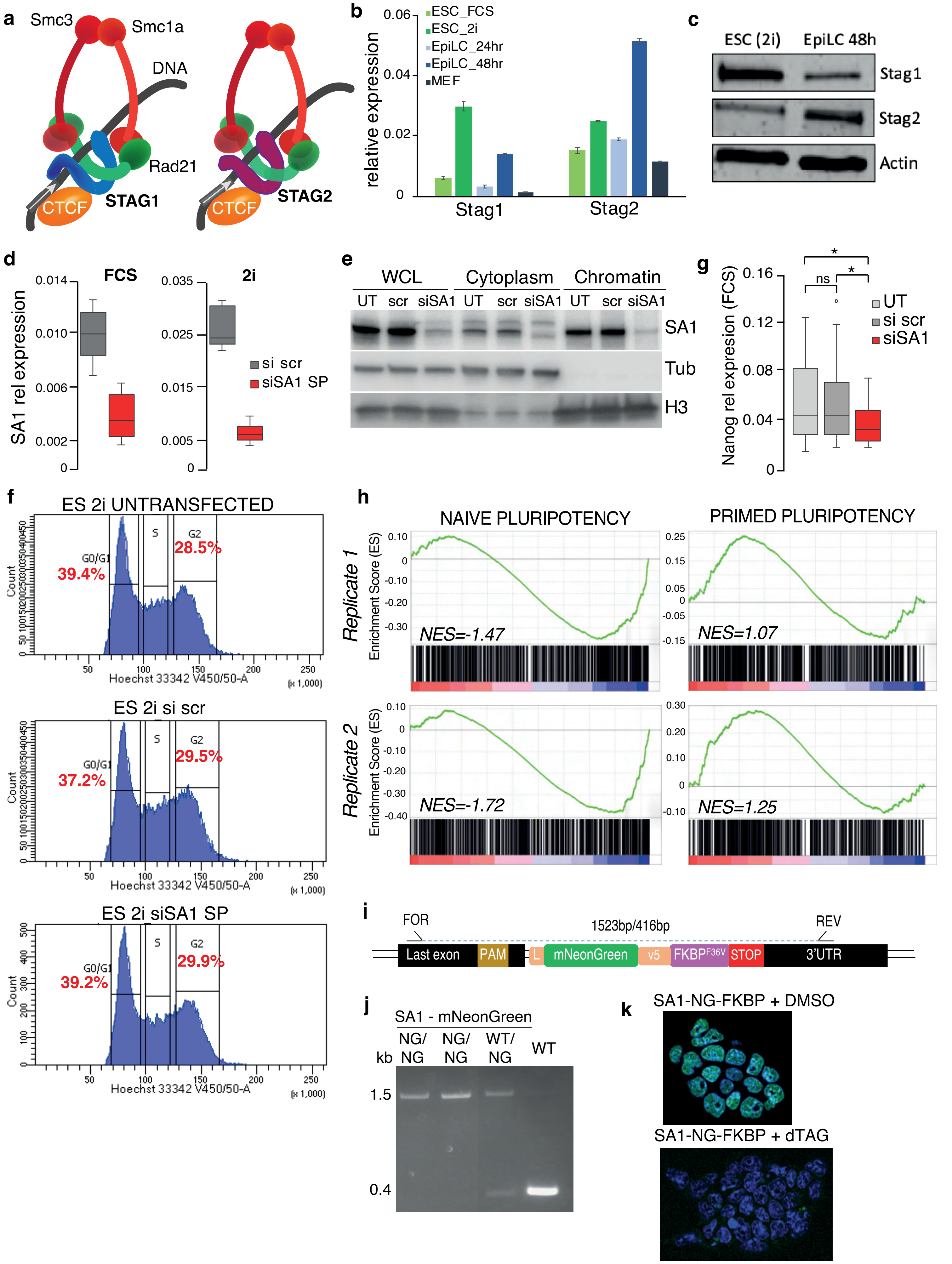

**Supplementary Figure 2.** *Pezic et al.*

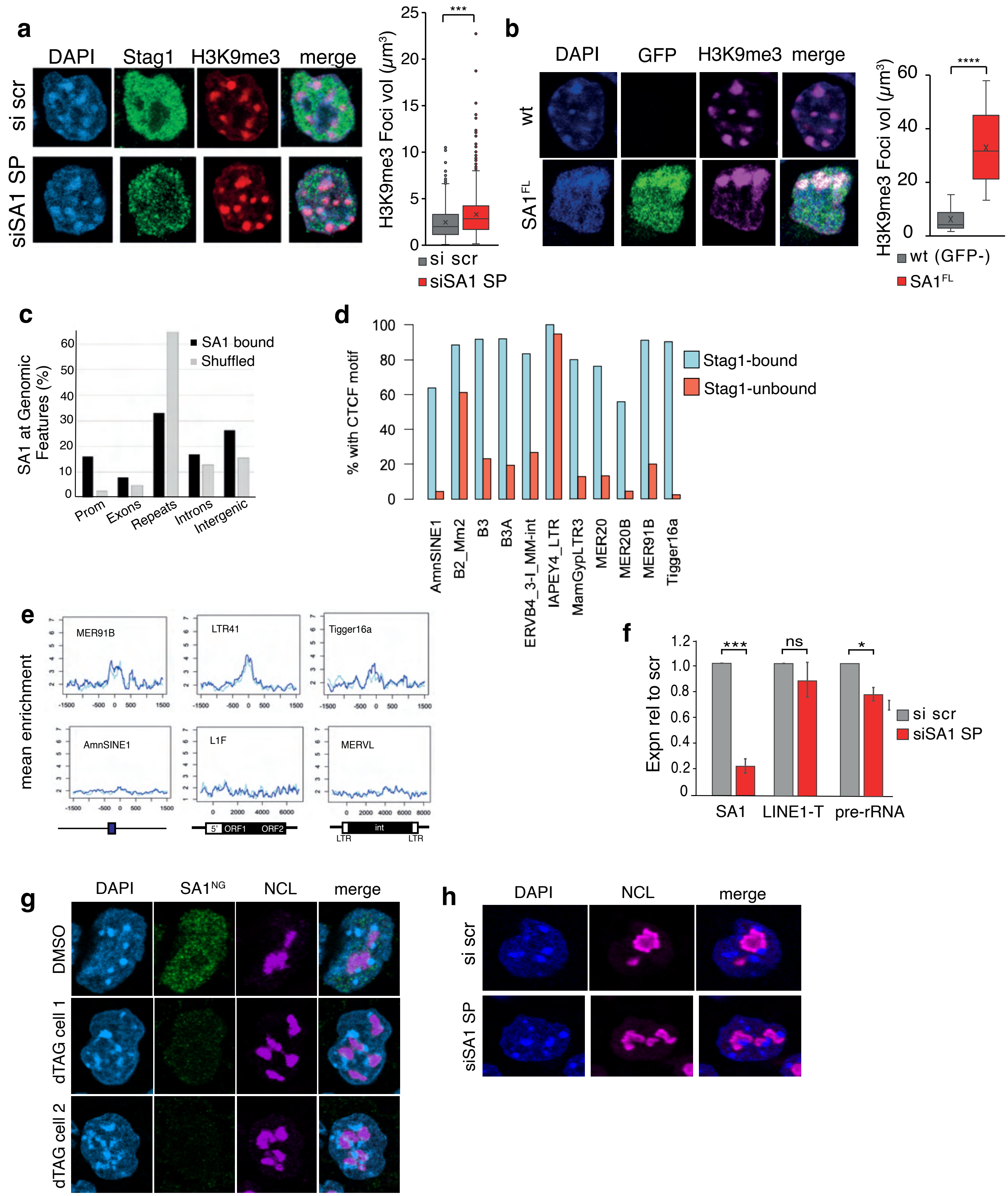

**Supplementary Figure 3.** *Pezic et al.*

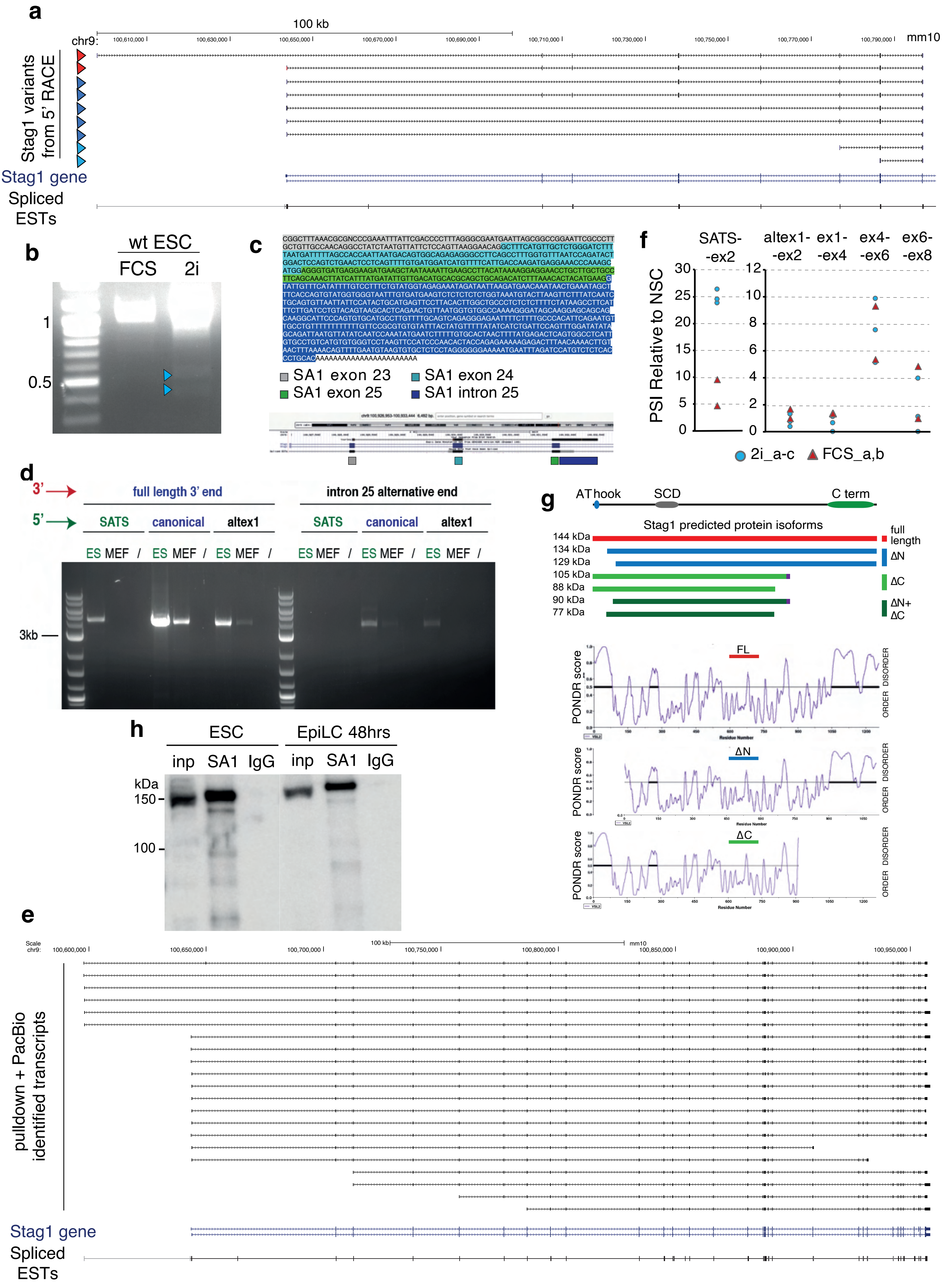

**Supplementary Figure 4.** *Pezic et al.*

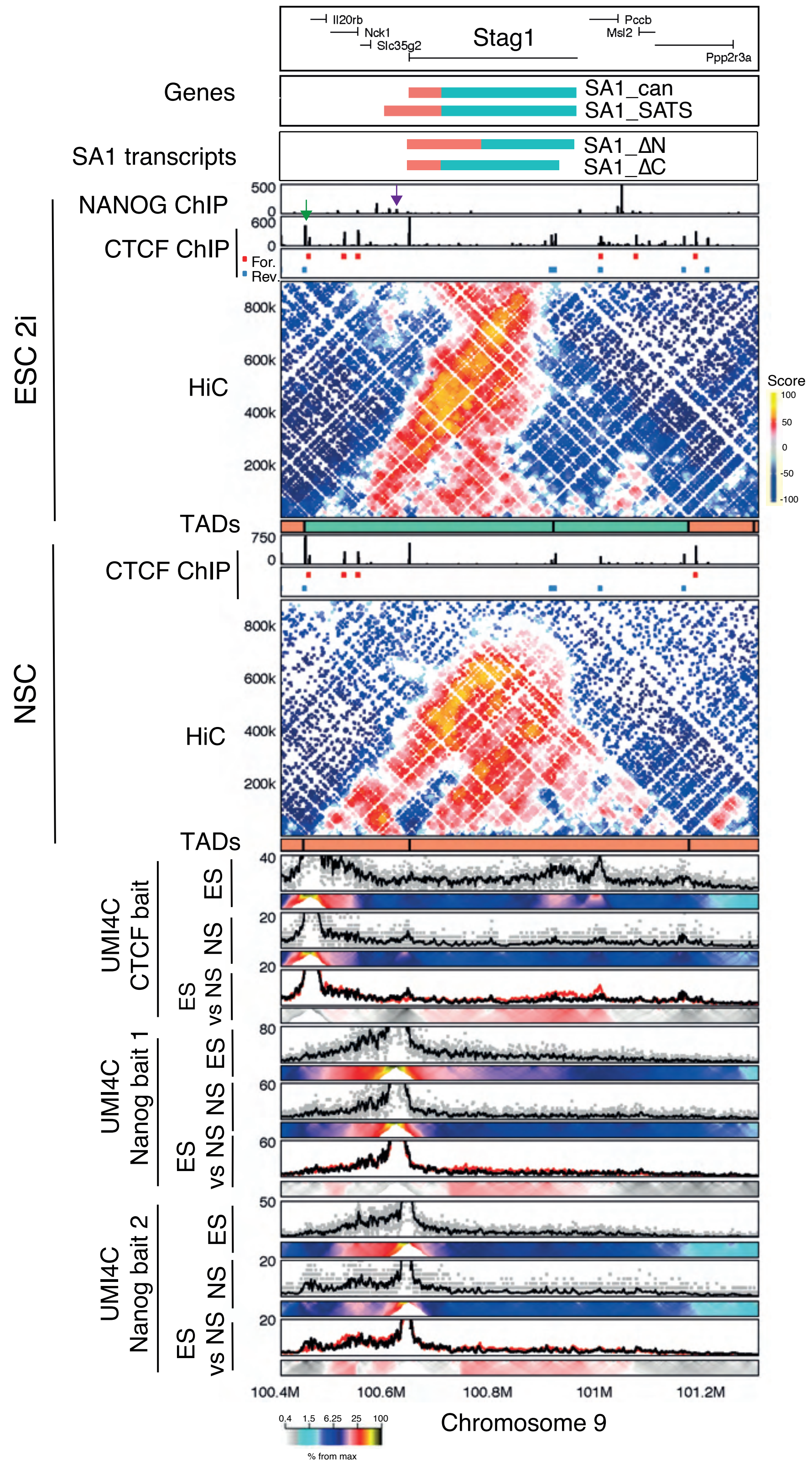

**Supplementary Figure 5.** *Pezic et al.*

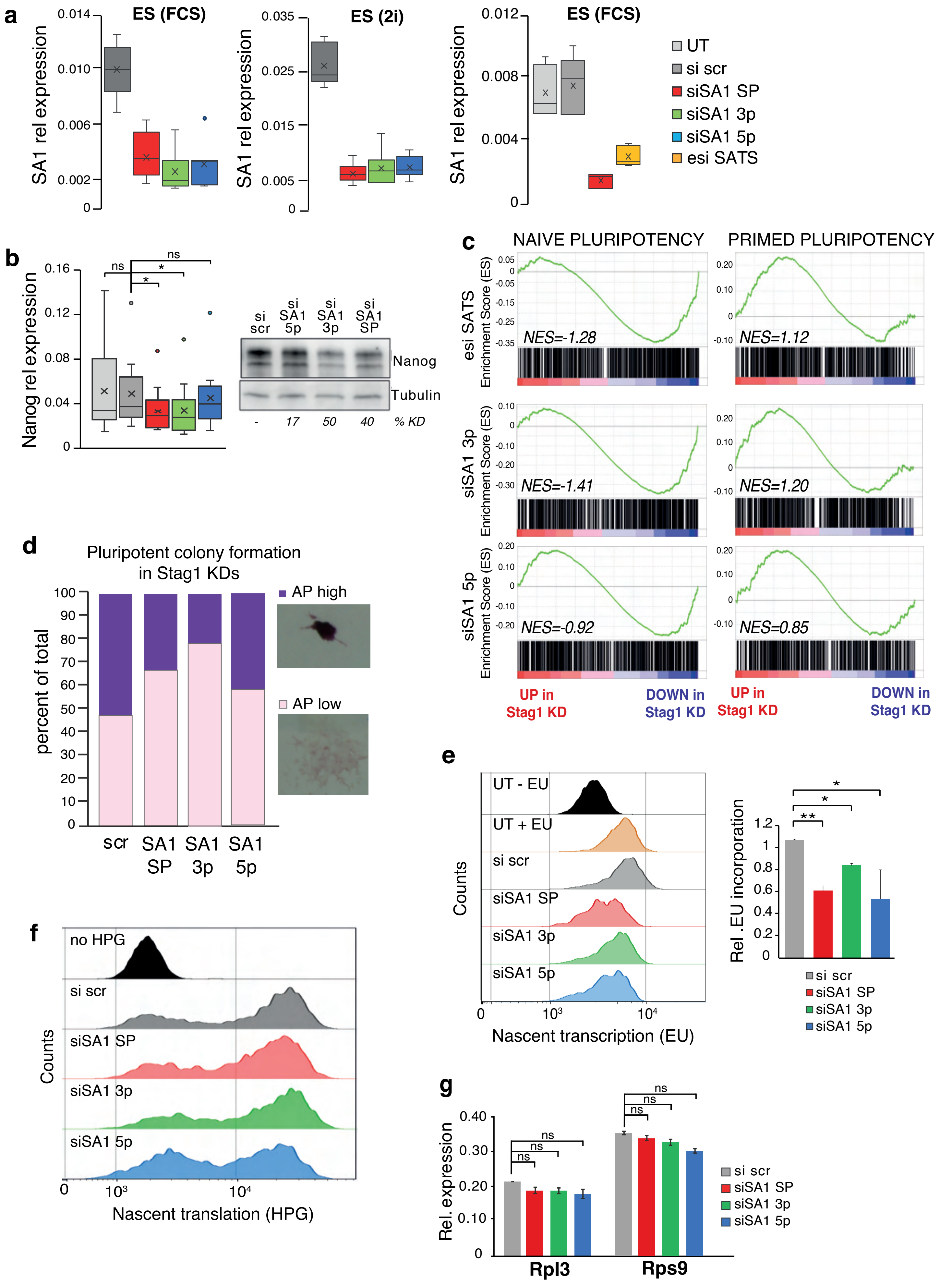

### Supplementary Figure 6.

*Pezic et al.*

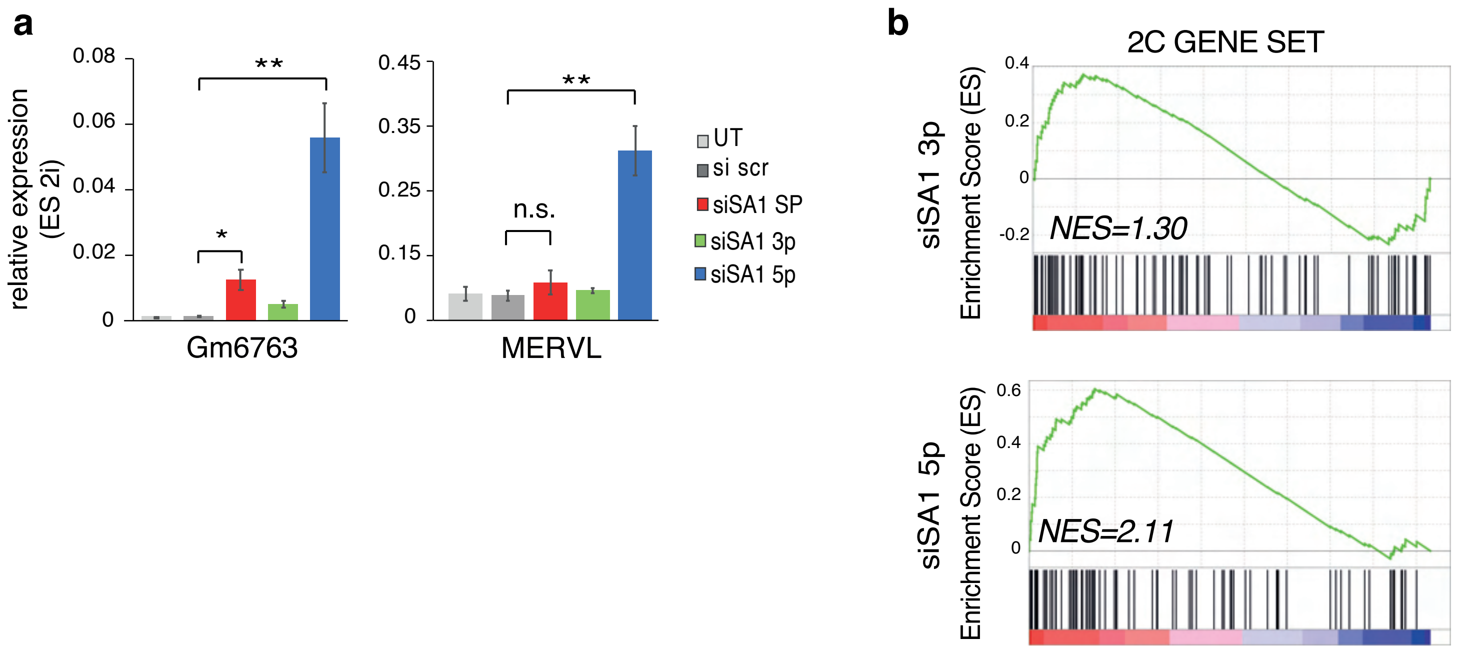
